# The mPFC molecular clock mediates the effects of sleep deprivation on depression-like behavior and regulates sleep consolidation and homeostasis

**DOI:** 10.1101/2025.06.09.658701

**Authors:** Wilf Gardner, David H. Sarrazin, Martin Balzinger, Carole Marchese, Axelle Ragno, Chockalingam Ramanathan, Maxime Veleanu, Stefan Vestring, Claus Normann, Patrice Bourgin, Tsvetan Serchov

## Abstract

Disruptions in sleep, circadian rhythms, and neural plasticity are closely linked to the pathophysiology and treatment of depression. Acute sleep deprivation (SD) produces rapid but transient antidepressant effects, yet the underlying mechanisms remain poorly understood. Using a mouse model of stress-induced depression, we found altered sleep architecture, impaired sleep homeostasis, and disrupted day-night oscillations of the markers of glutamatergic plasticity - Homer1a and synaptic AMPAR expression in the medial prefrontal cortex (mPFC). These changes were accompanied by a blunted homeostatic response to SD. We further show that SD and ketamine, both rapid-acting antidepressants, exert opposing effects on mPFC circadian gene expression: SD enhances the expression of negative clock loop genes (e.g., *Per*, *Cry*), mirroring stress effects, while ketamine downregulates these same genes. Targeted deletion of the core clock gene *Bmal1* in CaMK2a-expressing excitatory neurons of the mPFC disrupted sleep-wake architecture, elevated slow-wave activity, and abolished the behavioral and molecular (Homer1a) response to SD. Additionally, pharmacological activation of the clock repressor REV-ERB suppressed the antidepressant effects of SD. Our results demonstrate that the mPFC molecular clock is essential for the regulation of sleep consolidation and homeostasis, and mediates the effects of SD on behavior.

## INTRODUCTION

Major depressive disorder (MDD) is a major cause of disease-related reduction in quality of life worldwide and is among the most widespread mental health disorders (1). Standard first-line treatments, such as psychotherapy and monoamine-based antidepressants, often have significant drawbacks, including delayed onset of action, limited efficacy, and frequent side effects. Even with optimized treatment, only about half of patients achieve lasting remission, highlighting the need for more effective alternative therapeutic solutions (2). To address these challenges, chronotherapies have emerged as alternate treatment strategies (3), targeting sleep and circadian mechanisms, both of which are disrupted in depression (4).

Therapeutic sleep deprivation (SD) is a fast-acting antidepressant, with more than 50% of patients experiencing a significant reduction in depressive symptoms within hours. However, this effect is short-lived, with most responders relapsing after recovery sleep (RS) (5). Numerous studies have attempted to elucidate the mechanisms underlying SD, but the neurobiological basis of its antidepressant effects remains insufficiently understood (3, 6).

Impaired neural plasticity and disrupted information processing within neural networks have been suggested as key mechanisms underlying MDD, replacing previously dominant ideas relating primarily to monoamine activity (7–12). Moreover, a growing body of evidence indicates that the restoration of synaptic plasticity represents a key factor in the treatment response to rapid antidepressant interventions (13–17). Indeed, we have previously demonstrated that the induction of the synaptic protein Homer1a and concomitant modulation of glutamatergic signaling in the medial prefrontal cortex (mPFC)—a brain region heavily implicated in depression (18)—plays a key role in the antidepressant effects of SD and ketamine (19–23). Moreover, recent studies suggest that the molecular mechanisms of action of ketamine and SD involve the modulation of both synaptic plasticity and the circadian clock (6, 20, 24–27).

The circadian clock governs numerous biological and behavioral processes, including sleep, and is synchronized by the master circadian pacemaker in the suprachiasmatic nucleus (SCN). The molecular clock is regulated by feedback/feedforward loops driven by the transcription and translation of core clock genes. The positive regulators BMAL1 and CLOCK cyclically activate the transcription of suppressor genes Period (Per1, Per2) and Cryptochrome (Cry1, Cry2), whose proteins subsequently inhibit BMAL1/CLOCK-mediated activation. Additionally, a secondary regulatory loop involves retinoic acid receptor-related orphan receptors (ROR α, β, γ) and nuclear receptors REV-ERB (α, β), which act respectively as clock activators and repressors (28).

MDD is also associated with disruption of circadian rhythms (29) and sleep regulation, typically manifesting in increased sleep fragmentation, disinhibition of rapid eye movement (REM) sleep, and decreased slow-wave sleep (SWS) (4, 30). Reductions in the amplitude and changes to the distribution of high-amplitude low-frequency slow-wave activity (SWA) during SWS may be linked to impaired plasticity in depression. Indeed, the therapeutic effects of SD are correlated with the modulation of SWA/SWS (16, 17, 26, 31–34).

In this study, we initially examined sleep regulation and homeostatic glutamatergic plasticity in the mPFC of a mouse model of a stress-induced depression-like phenotype. Our findings reveal altered sleep architecture and homeostasis in these mice, alongside dysregulation of the day-night oscillating expression of Homer1a and synaptic AMPARs, two markers of glutamatergic plasticity. Additionally, they exhibit blunted response to SD, characterized by impaired rebound SWA, attenuated RS, and altered upregulation of Homer1a and AMPAR. Then, we investigated the involvement of mPFC molecular clock in sleep regulation and SD response. Our data indicate that ketamine and SD modulate the mPFC molecular circadian clock in distinct ways. 6h SD increased the expression of negative molecular clock regulators, lasting even after the RS period, while ketamine causes downregulation of the same clock suppressors. Using virally mediated *Bmal1*KO, we also show that the specific disruption of the circadian clockwork in the CaMK2a glutamatergic neurons of the mPFC dysregulates the homeostatic drive for sleep, enhances SWA, and impairs the therapeutic response to SD via inhibition of Homer1a induction. Finally, we demonstrate that pharmacological potentiation of the circadian clock suppressor REV-ERB blocks the rapid antidepressant effects of SD. Thus, our findings highlight the critical role of the mPFC molecular clock in sleep regulation and the response to SD, indicating a potential link between circadian and plasticity mechanisms.

## MATERIALS AND METHODS

### Animals

Adult wild type C57BL/6J (RRID: IMSR_JAX:000664) and *Bmal1* floxed mutant mice (B6.129S4(Cg)-*Arntl^tm1Weit^*/J; Jax Stock#: 007668; RRID: IMSR_JAX:007668) (35) were sourced from Charles River (France and Germany). All mice were at least 8 weeks old at the start of experimental procedures. Both male and female mice were utilized throughout the entirety of the study, with the sex of the animals specified for each experiment in Supplementary Table 1. The mice are housed in a temperature- and humidity-controlled environment, with *ad libitum* access to food and water, maintained on a 12-hour light-dark cycle. The light phase corresponds to the resting period, while the dark phase represents the active period for the mice. The time points of the behavioral assessments were aligned with the timing of mouse sacrifice and brain tissue collection for subsequent molecular analyses. The Chronobiotron animal facility is registered for animal experimentation (Agreement A67-2018-38). All procedures were performed in accordance with the German animal protection law (TierSchG), FELASA (http://www.felasa.eu), the guide for care and use of laboratory animals of the national animal welfare body GV-SOLAS http://www.gv-solas.de) and the EU Directive 2010/63/EU for animal experiments and were approved by the animal welfare committee of the Universities of Freiburg (X-20/06R, 35-9185.81/G-19/44 and G-15/117) and Strasbourg (CREMEAS, APAFIS n°2020042818477700) as well as by local authorities.

### Surgical procedures

All stereotaxic surgeries were conducted under zolazepam/xylazine intraperitoneal anesthesia (Zoletil 50, 40 mg/kg, Paxman, 10 mg/kg (Virbac, France)). Prior to incision, animals received analgesia via subcutaneous (s.c.) NSAID meloxicam (Metacam, 2mg/kg, (Boehringer Ingelheim, Germany)) and local anaesthetics lidocaïne and bupivacaïne at the incision site (Lurocaïne, (Vetoquinol, Canada)/Bupivacaïne (Pfizer, USA), 1 mg/kg). Ophthalmic gel (Ocrygel, TVM, UK) was applied to prevent corneal drying, and body temperature was maintained using a heated pad. Hydration was ensured during and following surgery with subcutaneous saline injections. Stereotactic coordinates were measured from bregma with the skull fixed in the flat head position, with dorsal-ventral measurements taken from the level of the dura. The mPFC was targeted using the following coordinates (in mm): anterior-posterior +1.70; medio-lateral ±0.35; dorso-ventral -2.20. After surgery, animals received atipamezole to reverse the effects of xylazine (Revertor (Chanelle Pharma, Ireland), 0.1 mg/kg, s.c.) and further meloxicam delivered via drinking water for three days (Inflacam (Virbac, France), 5 mg/kg). All animals were allowed a minimum 7-day recovery period before further experimental procedures.

### *In vivo* stereotaxic microinjections of recombinant adeno-associated viral vectors

In vivo injection of Cre- and EGFP (control) expressing viruses (pENN.AAV.CamKII.HI.GFP-Cre.WPRE.SV40 was a gift from James M. Wilson (Addgene viral prep #105551-AAV9; http://n2t.net/addgene:105551; RRID: Addgene_105551) and pAAV-CaMKIIa-EGFP was a gift from Bryan Roth (Addgene viral prep #50469-AAV9; http://n2t.net/addgene:50469; RRID: Addgene_50469) respectively). The AAVs were injected bilaterally into the mPFC via Hamilton syringe fitted with a 33-gauge needle, with a delivery rate set at 200 nl/min and a final volume of 650 nl/side. Following injection, the needle was briefly retained in place and then slowly withdrawn at a rate of 1 mm/min. Mice were allowed a 3-week recovery period post-injection to ensure sufficient viral expression. Virus transfection localization and knockdown efficacy were verified by immunohistochemistry and qRT-PCR.

### Drug treatment

Mice received an intraperitoneal (i.p.) injection of racemic ketamine (3 mg/kg ±ketamine hydrochloride, Sigma-Aldrich) dissolved in saline 0.9% NaCl (Aquapharm) or SR10067 (30mg/kg, Tocris) dissolved in Cremophor EL (Merck)/DMSO (Sigma)/saline 0.9% 15:10:75% by volume (36) at the indicated time before behavioral testing or sacrifice.

### Sleep deprivation (SD)

SD was performed using the gentle handling method that includes touching the animals with the hand or a brush, or gently shaking or tapping at the cage. The mice were sleep-deprived for 6h starting at the beginning of the light phase (ZT00). When indicated, SD was followed by recovery sleep (RS) period.

### Behavioral studies

During tail suspension and forced swim tests, the animal’s behavior was video recorded (Debut Professional v8.23 video capture software, NCH Software) and manually scored by two independent experimenters, both blinded to the experimental conditions. Tail suspension test, forced swim test and the chronic despair paradigm were performed at ZT06 (6h after lights on at ZT00) with both male and female mice. Activity of the mice were monitored using an IntelliCage system (TSE Systems). To avoid aggressive behaviors only female mice were used for the IntelliCage system experiments.

#### Tail suspension test (TST)

For TST, mice were attached by their tails (1–1.5 cm from the tip of the tail) to a horizontal bar. Each trial was conducted for 6 min and immobility time was recorded. Mice observed to climb their tails (> 10% of total time) were eliminated from further analysis.

#### Forced swim test (FST)

For the classical FST, mice were placed in a transparent glass cylinder (15 cm diameter) filled to a height of 20 cm with water (22-25°C). Immobility time was assessed during a 10 min swim session. Mice were considered to be immobile when they floated in an upright position and made only minimal movements to keep their head above the water.

#### Chronic behavioral despair model (CDM)

To induce chronic depression-like behavior, mice were subjected to daily swim sessions in a transparent glass cylinder (15 cm in diameter) filled with water (22-25°C) to a depth of 20 cm, for 10 minutes per day over 5 consecutive days (induction phase). To avoid potential circadian disruption, control mice were awakened and temporarily relocated to a new cage during the swim session of the CDM mice. After a one-week rest period, a final swim session was conducted (test phase). Repeated swim exposure resulted in a sustained increase in immobility time and a decrease in sucrose preference during the test phase. This method has been established as a model for depression-like behavior in mice and reliably predicts the effects of antidepressants (19, 21, 37–41).

#### Activity analysis in the IntelliCage

The IntelliCage system (TSE Systems) automated analysis of spontaneous exploratory behavior and activity patterns in up to 16 group-housed mice, each implanted with a radio-frequency identification (RFID) transponder. The unit consists of an open area with 4 red housings in the center and 4 recording corners. Mice have free access to food in the middle top of the IntelliCage, while water is available at the corners behind remote-controlled guillotine doors. Each corner contains 2 drinking bottles and can accommodate only one mouse at a time. Behavioral parameters, including the number and duration of visits to the corners, nosepokes toward the doors, and licking of the bottles, were monitored using PC-based tracking software (IntelliCage Plus, TSE Systems). Prior to testing, the mice underwent a minimum 5-day adaptation period, during which water was available *ad libitum* in all corners. The circadian locomotor and exploratory activity of the mice measured as corner visits was analysed using the ClockLab software (Actimetrics).

### *In vivo* electrophysiology recordings

Two electrocorticogram (ECoG) electrodes (constructed from 1.2 mm gold-plated steel screws and 60µm-diameter Teflon-coated tungsten wire (World Precision Instruments, USA)), were implanted at the level of the dura over the frontal and parietal regions. Reference and ground screw electrodes were also inserted, and an additional electrode (60µm-diameter Teflon-coated tungsten wire (World Precision Instruments, USA)) was placed in the nuchal muscle to record electromyogram (EMG) activity. The electrodes were secured to the skull with Superbond (Sun Medical), connected to a connecting headpiece (P1 Technologies), and fixed with dental cement. Following recovery from surgery, animals were attached *via* a flexible cable to a rotating joint (P1 Technologies), allowing free movement within their home cage. The animals were habituated to the recording setup for 72 hours prior to any recording procedures. During recording sessions, electrophysiological signals were amplified, digitized, and sampled at 500 Hz (Neuvo 64-channel amplifier and ProFusion software package, Compumedics, Australia), and data stored for offline analysis.

### Sleep scoring and spectral analysis

Analysis of vigilance states was performed manually using ProFusion software (Compumedics, Australia). ECoG and EMG signal were segmented into 4-second epochs, with each epoch scored according to standard criteria into one of three states: wake (identified by ECoG activity in the theta (8-12Hz) and high frequency ranges, with concurrent EMG activity), SWS (identified by characteristic slow waves of low-frequency, high-amplitude ECoG activity and low EMG), or REM sleep (identified by characteristic theta and minimal EMG activity). After scoring, sleep data parameters were averaged in 1-hour intervals.

ECoG signal was imported into MATLAB (Mathworks, USA) for spectral analysis using the Chronux (Bokil et al., 2010) and MATLAB Signal Processing (Mathworks, USA) toolboxes. Line noise was removed using a 512-point Hanning window, and values exceeding 3 standard deviations of the raw data identified as artefacts and removed with 1-second adjacent data. The signals were separated according to vigilance state (as previously described), concatenated, bandpass filtered between 0.5 and 200 Hz and transformed using a multi-taper method (time-bandwidth product of 3, using 5 Slepian tapers with a 2-second window moving at 0.1 second) implemented via the Chronux toolbox. Data were z-transformed and averaged over 2h periods prior to statistical analysis.

### Quantitative real-time PCR (qRT-PCR)

Mice from each experimental group were killed by cervical dislocation. The brains were rapidly extracted, coronally sectioned, and the mPFC and SCN regions were microdissected, quickly frozen on dry ice and stored at -80°C until used for RNA isolation. The RNA extraction and expression analyses were performed as previously described (21). Briefly, the tissues were homogenized in guanidine thiocyanate/ 2-mercaptoethanol buffer and total RNA was extracted using the sodium acetate/phenol/chloroform/isoamyl alcohol method. Then, samples were isopropanol-precipitated and washed twice with 70% ethanol. The resulting pellets were dissolved in RNase-free Tris-HCl buffer (pH 7.0), and RNA concentrations were measured using a spectrophotometer (BioPhotometer; Eppendorf). Reverse transcription was performed with 1 mg of total RNA using M-MLV reverse transcriptase (Promega). Quantitative real-time PCR was performed on a 7300 Real-Time PCR System with Sequence Detection Software qPCR v1.3.1 (Applied Biosystems), utilizing Fast SYBR^TM^ Green Master Mix (4385612, Applied Biosystems). All qRT-PCR experiments were performed blinded as the coded cDNA samples were pipetted by a technician. The target gene mRNA levels were normalized to the levels of apolipoprotein E (*ApoE*), glyceraldehydes-3-phosphate dehydrogenase (*GAPDH*) and *s12* RNA via geometric averaging of multiple internal control genes (42). The primer sequences used were as follows - *ApoE*: 5‘-CCTGAACCGCTTCTGGGATT-3‘, 5‘-GCTCTTCCTGGACCTGGTCA-3‘; *GAPDH*: 5‘-TGTCCGTCGTGGATCTGAC-3‘, 5‘-CCTGCTTCACCACCTTCTTG-3‘; *s12*: 5‘-GCCCTCATCCACGATGGCCT-3‘, 5‘-ACAGATGGGCTTGGCGCTTGT-3‘; *Per1*: 5’-CCAGATTGGTGGAGGTTACTGAGT-3’, 5’-GCGAGAGTCTTCTTGGAGCAGTAG-3’; *Per2*: 5’-AGAACGCGGATATGTTTGCTG-3’, 5’-ATCTAAGCCGCTGCACACACT-3’; *Cry1*: 5’-AGGAGGACAGATCCCAATGGA-3’, 5’-GCAACCTTCTGGATGCCTTCT-3’; *Cry2*: 5’-GCTGGAAGCAGCCGAGGAACC-3’, 5’-GGGCTTTGCTCACGGAGCGA-3’; *Bmal1*: 5’-CTCCAGGAGGCAAGAAGATTC-3’, 5’-ATAGTCCAGTGGAAGGAATG-3’; *Clock*: 5’-GGCGTTGTTGATTGGACTAGG-3’, 5’-GAATGGAGTCTCCAACACCCA-3’; *Rev-erbα*: 5’-CCCTGGACTCCAATAACAACACA-3’, 5’-GCCATTGGAGCTGTCACTGTAG-3’; *Rorα*: 5’-TTGCCAAACGCATTGATGG-3’, 5’-TTCTGAGAGTCAAAGGCACGG-3’; 5’-TGTAAGTGTGTCTGCTCCGCG-3’; Homer1a: 5‘-CAAACACTGTTTATGGACTG-3‘, 5‘-TGCTGAATTGAATGTGTACC-3‘.

### Preparation of synaptosomal extracts

Mice were killed by cervical dislocation, and the dissected mPFC regions were mechanically homogenized in homogenization buffer (320 mM sucrose, 4 mM HEPES [pH 7.4], 2 mM EDTA) and centrifuged at 800 g for 15 min at 4°C to obtain total (S1) and nuclear fraction (P1). All buffers contained phosphatase (P5726-1ML) and protease inhibitor (P8340-1ML) cocktail (Sigma-Aldrich). Synaptic (P2) and cytosolic (S2) fractions were obtained by centrifugation of S1 at 10 000 g for 15 min at 4°C. Washed synaptosomal pellets (P2) were directly lysed in lysis buffer (50 mM Tris-HCl [pH 6.8], 1.3% SDS, 6.5%, glycerol, 100 mM sodium orthovanadate) containing phosphatase and protease inhibitor cocktail (Sigma-Aldrich), then boiled for 5 min at 95°C before processing for SDS-PAGE and western blot. The protein concentration of the different fractions was determined using a BCA assay kit (Thermo Fisher Scientific) according to the manufacturer’s instructions.

### Western blot

After the addition of 10 mM DTT, synaptosomal (P2) lysates were boiled for 5 min at 95°C, separated by SDS-PAGE on 7.5% acrylamide gels and transferred to nitrocellulose transfer membrane (Whatman). The membranes were blocked with 5% non-fat dry milk in TBS-T (1% Tween20 in Tris-buffered saline (TBS)) and afterward incubated with the respective primary antibodies diluted in TBS (mouse anti-GluR1-NT, Merck (MAB2263), 1:1000; rabbit anti-Homer1, Merck (ABN37), overnight at 4°C. After 3 washes, membranes were incubated for 1 hour at room temperature with horseradish peroxidase-conjugated secondary antibodies diluted in TBS-T: sheep anti-mouse (GE Healthcare, NA931, 1:20,000) and donkey anti-rabbit (GE Healthcare, NA9340, 1:25,000). The membranes were then developed using an Amersham Imager 680 (GE Healthcare) using enhanced chemiluminescence detection kit (Thermo Fisher Scientific). Band intensities were quantified by densitometry with ImageJ 1.53m software (National Institute of Health, USA) and normalized to the appropriate loading control.

### Immunofluorescence

Animals were anesthetized using a mix of Ketamin-Rompun (Ketamine [CP Pharma] 50 mg and Rompun [Bayer Healthcare] 0.5 mg per 100 g body weight) and transcardially perfused with 50 mL of ice-cold PBS (8.1 mM Na2HPO4, 138 mM NaCl, 2.7 mM KCl and 1.47 mM KH2PO4 [pH 7.4]). The brains were then extracted, postfixed overnight in 4% paraformaldehyde in PBS at 4°C and cryoprotected for 2 days in 30% sucrose in PBS at 4°C. Brains were then frozen and 40 μm coronal sections were cut with a sliding cryostat (Leica Microsystems). Slices were washed in PBS and mounted on slides using mounting medium from Life technologies (E6604). All fluorescence images were detected and photographed with an AXIO Imager.M2 fluorescence microscope and analyzed using ZEN 3.5 software (Carl Zeiss) and ImageJ 1.53m (National Institute of Health, USA).

### Statistical analysis

All values are presented as means ± SEM. Statistical analyses were performed with GraphPad Prism 8.0.0 software (GraphPad Software) using one- or two-way analysis of variance (ANOVA) followed by Bonferroni’s or Tukey’s post hoc tests for comparisons between multiple groups, or an unpaired two-tailed Student’s t-test for comparisons between two groups. The daily rhythm of SWA was evaluated by cosinor analysis. Sine wave (least-squares regression) with the frequency constrained to exactly 24h was then fitted using the following equation: y = A + B*cos(2π [x − C]/24) where A is the mesor, B is the amplitude, and C is the acrophase of the fitted rhythm. To further compare the mesor, the amplitude and the acrophase of the curve fits generated from the sine wave model of the different treatment conditions the extra sum of squares F test was performed. A *P* value ≤ 0.05 was considered to be significant (\**P* ≤ 0.05, \*\**P* ≤ 0.01, \*\*\**P* ≤ 0.001). Prior to statistical analyses data assumptions (for example normality and homoscedasticity of the distributions) were verified using D’Agostino-Pearson, Shapiro-Wilk and Kolmogorov-Smirnov tests (GraphPad Prism 8.0.0 software). Detailed statistical approaches and results are provided in Supplementary Data Table 1. Statistical analysis summaries are mentioned in the figure legends. For all molecular and behavioral studies mice were randomly assigned to the groups. Most behavioral and molecular results were confirmed via independent replication of the experiments as shown in Supplementary Data Table 1. Additionally, investigators were blinded to the treatment group until data have been collected. Sample sizes were determined on the basis of extensive laboratory experience (19–21, 38) and were verified via power analysis.

### Reporting summary

Further information on research design is available in the Nature Research Reporting Summary linked to this article.

### Data and code availability

The data sets generated and/or analyzed during the current study can be found in the paper and the supplementary materials. Research materials will be made available on a reasonable request at.

## RESULTS

### Repetitive swim stress increases sleep fragmentation and SWS duration

To investigate the effects of a stress-derived depression-like phenotype on sleep, mice were subjected to the chronic behavioral despair paradigm (CDM), which induces passive immobility and anhedonia through daily 10 min swim sessions for 5 consecutive days (induction phase) (19–21, 37–41). CDM mice and naïve non-stressed controls were implanted with ECoG and EMG electrodes for continuous recording over 24h (Fig. 1a). Our data show that CDM mice exhibited several abnormalities in sleep architecture compared to naïve controls. Swim-stressed mice spent more time in SWS (Fig. 1c) at the expense of wake (Fig. 1b), reflected in increased total SWS time over the 24h and across both 12h dark and light cycles. In contrast, overall time spent in REM sleep was not affected (Fig. 1d). Moreover, CDM mice showed decreased locomotor and exploratory behavior during the first half of the active period (ZT12-18) and significantly delayed activity onset at ZT12, measured as corner visits in the IntelliCage (Supplementary Figs. S1a-c). The SWS of CDM mice was marked by increased fragmentation, with significantly more frequent transitions between wake and SWS than controls, particularly in the sleep-dominated light period (ZT12-24) (Fig. 1e). The average lengths of wake and SWS bouts were significantly shorter over the whole 24h and during the light period, respectively (Figs. 1f, g), reflecting the greater sleep fragmentation of SWS (Supplementary Figs. S1d-e). CDM mice entered REM sleep more frequently during the light period than controls (Fig. 1h), despite spending equal time in REM overall (Supplementary Figs. S1f, g).

**Fig. 1:**
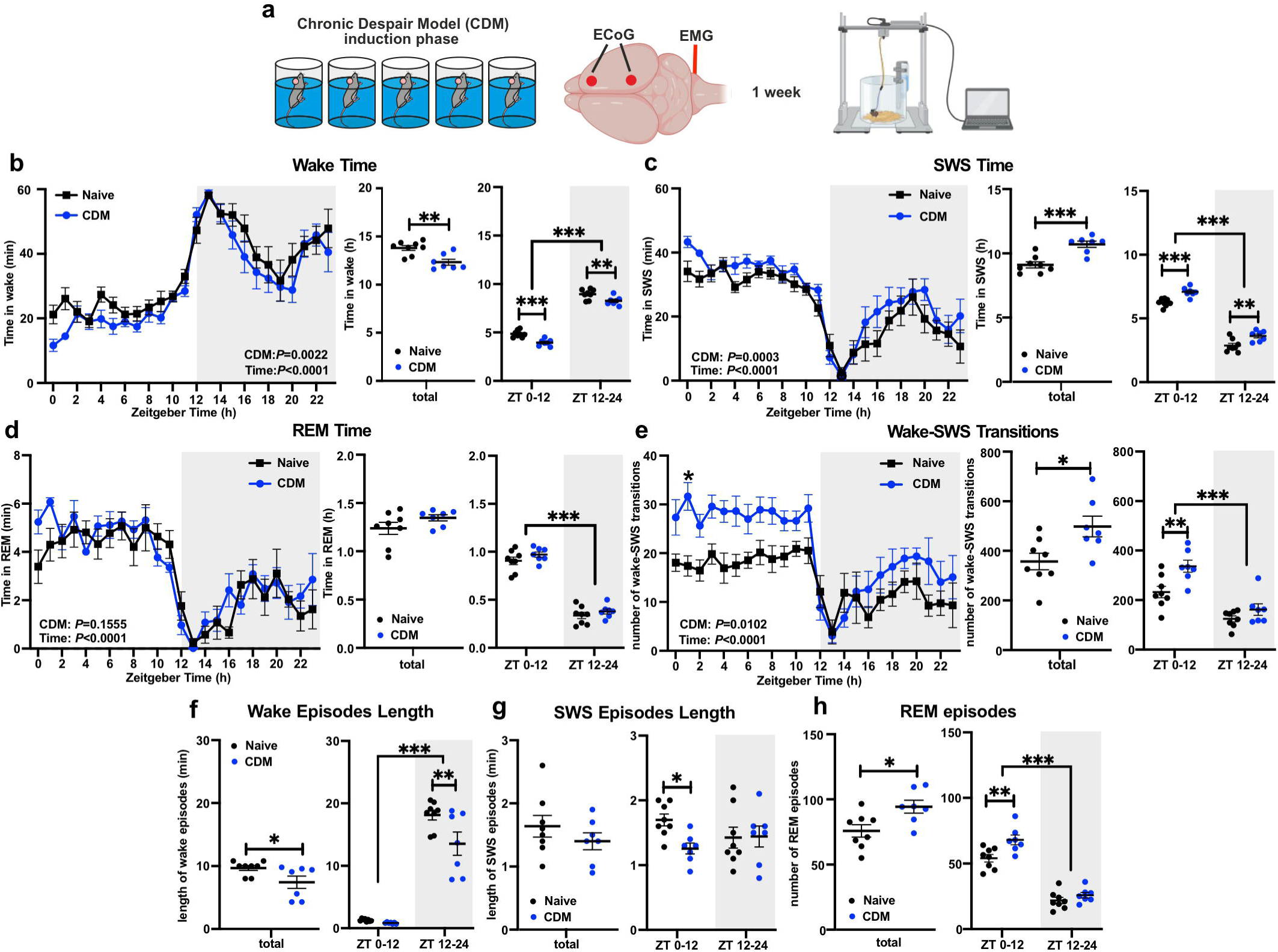
Repetitive swim stress increases SWS duration and sleep fragmentation. **a** Experimental design: WT mice were subjected to the CDM protocol before undergoing implantation of ECoG and EMG recording electrodes. After recovery, mice were recorded in their home cage. **b-h** Sleep architecture measures of naïve (control) (n=8) and CDM (n=7) mice over 24h. Time spent in wake (**b**), SWS (**c**) and REM sleep (**d**) per 1h periods over 24h (left), total time for 24h (middle) and time across 12h light (ZT00-12) and dark (ZT12-24) cycles (right). **e** Number of wake-SWS transitions per 1h periods over 24h (left), total number of transitions for 24h (middle) and across 12h light (ZT00-12) and dark (ZT12-24) cycles (right). Mean duration of spontaneous wake (**f**) and SWS (**g**) episodes for the whole 24h (left) and across 12h light (ZT00-12) and dark (ZT12-24) cycles (right). **h** Number of REM episodes for the whole 24h (left) and across 12h light (ZT00-12) and dark (ZT12-24) cycles (right). (**b-e** left & right plots and **f-h** right plots: repeated measures two-way ANOVA with Bonferroni post-hoc test; **b-e** middle plots & **f-h** left plots: two-tailed Student’s t-test; \**P*<0.05, \*\**P*<0.01, \*\*\**P*<0.001). Data are presented as mean ±SEM and the individual data points are depicted. See also Supplementary Fig. S1 and Supplementary Data Table 1. Some of the sketches were made with biorender.com

Taken together these data indicate that the repeated-swim stress in CDM paradigm alters sleep architecture causing enhanced SWS duration and sleep fragmentation, marked by increased wake-SWS transitions, shortened SWS and wake bouts, and higher number of REM sleep episodes.

### Dysregulated sleep homeostasis and homeostatic plasticity, and altered response to sleep deprivation in CDM animals

Next, we examined CDM effects on sleep homeostasis and homeostatic plasticity. SWA during SWS is associated with both homeostatic drive for sleep and crucial mechanisms linking to neuronal plasticity, such as synaptic downscaling (32, 33, 43–45). Our data show that the CDM did not significantly change the delta power of the SWS. However, the rhythmic pattern of SWA distribution across 24h was affected in the CDM mice, leading to marked flattening of the fitted sine wave curve and robust rhythm amplitude decrease (Fig. 2a, Supplementary Fig. S2a). Moreover, while delta power is significantly reduced during the sleep-dominated light period (ZT00 vs ZT12) in naïve controls, the CDM mice showed no significant change in SWA (Figs. 2b, Supplementary Fig. S2b).

**Fig. 2:**
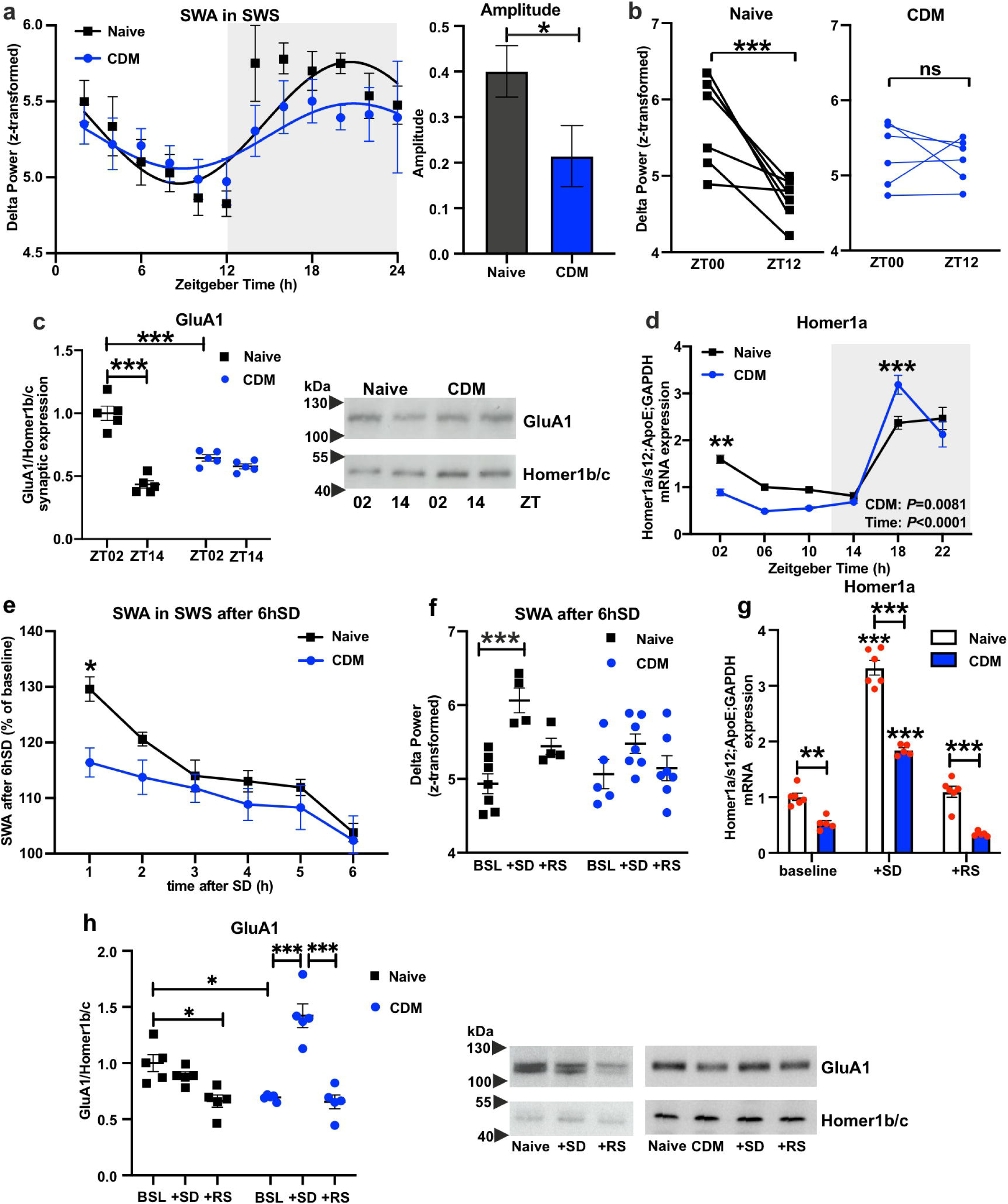
Dysregulated sleep homeostasis and homeostatic plasticity, and altered response to sleep deprivation in CDM animals. **a** Slow wave activity (SWA) presented as delta power (0.5-4Hz) of ECoG signal recorded during slow wave sleep (SWS) per 2h episodes across 12h:12h LD conditions including fitted sine waves (left) and the calculated amplitude of rhythmic oscillation (right) of naïve (n=8) and CDM (n=6) mice (left: repeated measures mixed-effects model two-way ANOVA, nonlinear regression sine wave fit and cosinor analyses; right: extra sum of squares F test: \**P*<0.05). **b** SWA in SWS at the start (ZT00) and at the end (ZT12) of the sleep period of naïve (n=8) and CDM (n=6) mice (repeated measures two-way ANOVA with Bonferroni post-hoc test: \*\*\**P*<0.001). **c** Quantitative data (left) and representative western blots (right) of synaptic AMPA receptors GluA1 subunit expression normalized to Homer1b/c in mPFC of naïve and CDM mice at ZT02 and ZT14. (n=5, two-way ANOVA with Bonferroni post-hoc test: \*\*\**P*<0.001). **d** Relative mRNA expression of Homer1a in mPFC tissue harvested from naïve (control) and CDM mice every 4h at 12:12h LD condition (n=5 mice per group, one-way ANOVA with Bonferroni post-hoc test: \*\**P*<0.01, \*\*\**P*<0.001 naive vs. CDM). **e** Time course of SWA during SWS after 6h of acute sleep deprivation (6hSD) in percentage relative to the baseline average of naïve (n=4) and CDM (n=6) mice (repeated measures two-way ANOVA with Bonferroni post-hoc test: \**P*<0.05). **f** SWA during SWS at ZT06 at baseline (BSL) after 6hSD and following 24h of recovery sleep (RS) of naïve (n=4) and CDM (n=6) mice (repeated measures two-way ANOVA with Bonferroni post-hoc test: \*\*\**P*<0.001). **g** Relative mRNA expression of Homer1a in mPFC at ZT06 of naive (n=6) and CDM (n=5) mice at baseline, after 6hSD and following 24h of recovery sleep (RS) (two-way ANOVA with Bonferroni post-hoc test: \*\**P*<0.01, \*\*\**P*<0.001 in comparison to baseline). **h** Quantitative data (left) and representative western blots (right) of synaptic GluA1 AMPA receptors expression normalized to Homer1b/c in mPFC of naïve and CDM mice at ZT06 BSL, after 6hSD and following 24hRS (n=5, two-way ANOVA with Bonferroni post-hoc test: \**P*<0.05, \*\*\**P*<0.001). Data are presented as mean ±SEM and the individual data points are depicted. See also Supplementary Fig. S2 and Supplementary Data Table 1.

The induction of the postsynaptic scaffolding protein Homer1a and the dynamic expression of synaptic AMPAR have been recognized as potent markers of homeostatic plasticity and correlated with SWA during SWS (45–47). Reflecting the SWA, naïve mice exhibited a ZT-dependent change in AMPA GluA1 synaptic expression in the mPFC between the early light period (ZT02) and early dark period (ZT14). However, the synaptic GluA1 expression in CDM was not different between ZT periods, and was significantly lower than controls at ZT02 (Fig. 2c). Likewise, oscillating expression of Homer1a was altered in the mPFC of stressed mice displaying lower levels at ZT02 and upregulated expression at ZT18 in comparison to controls with concomitant significantly increased amplitude and altered acrophase (Fig. 2d, Supplementary Fig. 2g).

In order to investigate SWA and homeostatic plasticity as functions of the homeostatic drive for sleep, CDM and naive mice were subjected to acute 6h SD (ZT00-ZT06). It has been previously proposed that the mechanism of action of therapeutic SD correlates with enhanced SWS and SWA, and it depends on Homer1a induction and synaptic AMPAR expression in the mPFC (16, 17, 19, 21, 34). In CDM mice, the SWA rebound following SD was significantly lower than that of control mice (Fig. 2e, Supplementary Fig. S2f), despite similar, but shorter, rebounds in SWS time (Supplementary Fig. S2c). While naive mice exhibited transiently elevated SWA relative to the equivalent ZT at baseline, the rebound seen in CDM was not significantly higher (Fig. 2f). Interestingly, acute SD caused an improvement in sleep fragmentation of CDM mice, reducing the wake-SWS transitions and increasing SWS episode length (Supplementary Figs. S2d, e). When examining Homer1a induction in the mPFC of CDM and naive mice, both groups showed a significant increase in Homer1a expression after 6hSD and a subsequent decrease in the following 24h recovery sleep (RS) period; however, CDM mice displayed lower expression at each condition compared to naive animals (Fig. 2g). In addition, CDM mice exhibited strong upregulation of synaptic GluA1 expression in mPFC as an immediate response to SD. In control mice, no such elevation was observed, with a significant reduction in expression from baseline seen after RS (Fig. 2h).

In summary, the CDM depression-like phenotype model exhibited altered SWA rhythm across 24h, correlating with disrupted regulation of the homeostatic plasticity (day-night dynamics of Homer1a and synaptic AMPAR expression) in the mPFC. Moreover, SWA, Homer1a, and AMPAR induction displayed differential response to SD in naïve and stressed mice.

### Rapid antidepressants — acute sleep deprivation and ketamine differentially affect mPFC clock gene expression

Numerous clinical studies have demonstrated that acute SD and subanesthetic dose of ketamine produce both rapid and robust antidepressant effects in depressed patients; however, ketamine’s action is longer-lasting, whereas the therapeutic impact of SD relapses after a night of recovery sleep (5, 48). We investigated these effects in our CDM model and thus initially assessed the magnitude of action of ketamine or SD on the CDM mice (Fig. 3a). 6hSD (ZT00-ZT06) or ketamine (3mg/kg) (administered at ZT00) produced a rapid antidepressant response in the classical depression-like behavior tests FST and TST 6h following treatment (ZT06); this effect was maintained in ketamine-treated animals 24h later, while SD-treated animals reverted to a depressive-like phenotype after 24h RS (Figs 3b, c).

**Fig. 3:**
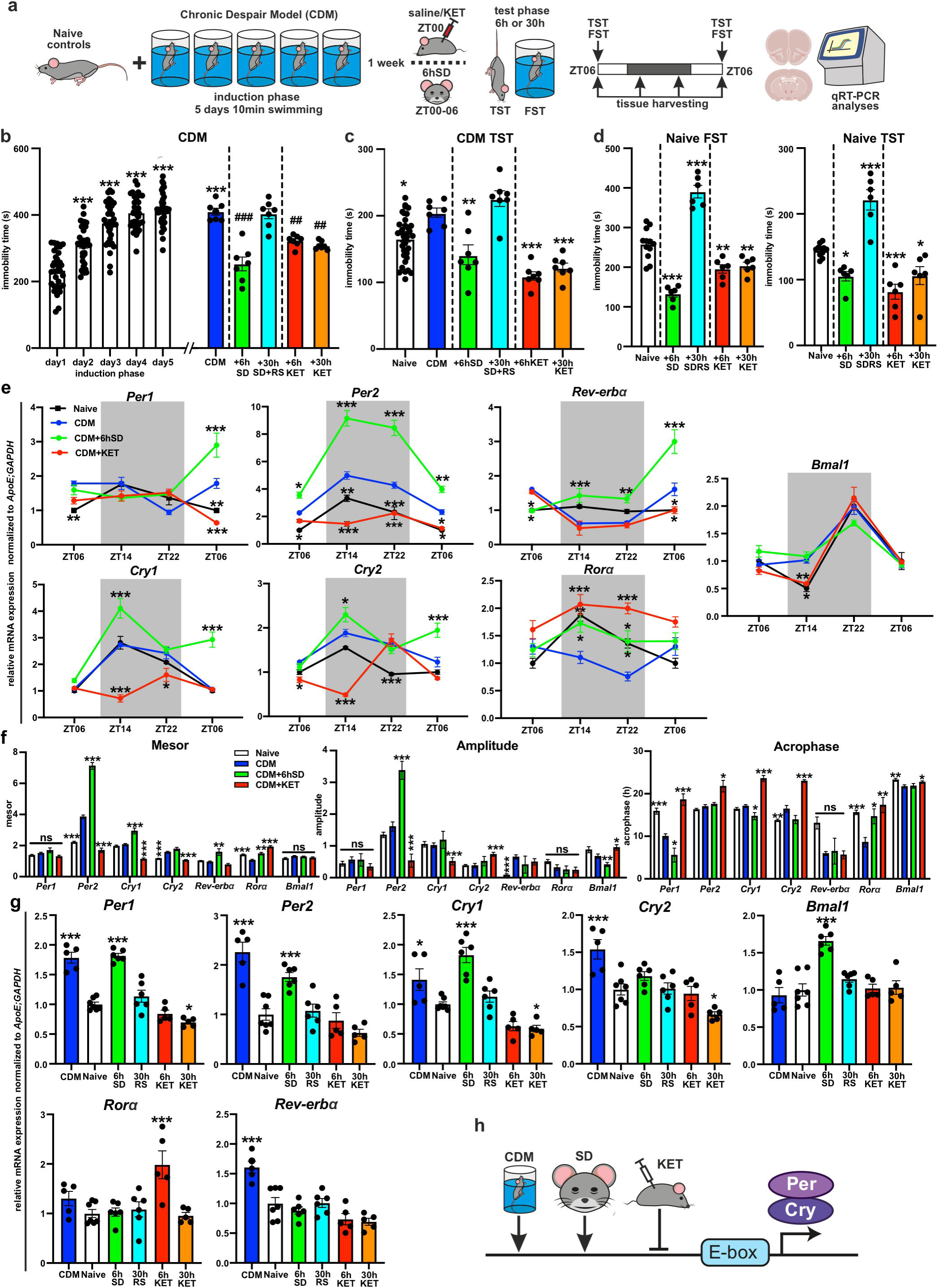
Rapid antidepressants acute sleep deprivation and ketamine differentially affect mPFC clock gene expression. **a** Schematic overview of the experimental design: naïve mice and mice subjected to the CDM paradigm were 1 week later divided into the following groups: naïve/CDM – saline i.p. injected at ZT00, +6hSD - sleep deprived for 6h (ZT00-ZT06), +30hSD+RS - sleep deprived for 6h followed by 24h recovery sleep, or i.p. injected with 3 mg/kg ketamine (KET) at ZT00; behavioral tests (TST and FST) were performed 6h or 30h after the start of the treatment (ketamine injection or SD) at ZT06; separate groups of animals were used for brain tissue harvesting (at the indicated time points) and gene expression analyses. **b** Immobility time spent during the induction (n=35) and test phase (n=7 per group) of CDM (repeated measures one-way ANOVA with Bonferroni post-hoc test: \*\*\**P*<0.001 in comparison to day1; ##*P*<0.01 and ###*P*<0.001 in comparison to the CDM group). **c** Immobility time in TST of naïve mice (n=35) and during the test phase of CDM (n=7 per group; repeated measures one-way ANOVA with Bonferroni post-hoc test: \**P*<0.05, \*\**P*<0.01, \*\*\**P*<0.001 in comparison to the CDM group). **d** Immobility time of FST (left) and TST (right) of naïve mice treated with 6hSD or ketamine (n=6 mice, one-way ANOVA with Bonferroni post-hoc test: \**P*<0.05, \*\**P*<0.01, \*\*\**P*<0.001 in comparison to the naïve group). **e** Relative mRNA expression of clock genes *Per1, Per2, Cry1, Cry2, Bmal1, Rorα* and *Rev-erb*α normalized to *ApoE* and *GAPDH* in mPFC samples from naïve or CDM mice injected at ZT00 with saline (CDM) or ketamine (KET) or 6hSD treated and harvested every 8h from ZT06 (end of SD or 6h after injection) till ZT06 (24h RS or 30h post injection) (n=5 mice per group, two-way ANOVA with Bonferroni post-hoc test: \**P*<0.05, \*\**P*<0.01, \*\*\**P*<0.001 vs. CDM). **f** Mesor, amplitude and acrophase of rhythmic expression (determined via nonlinear regression sine wave fit and cosinor analyses; extra sum of squares F test: \**P*<0.05, \*\**P*<0.01, \*\*\**P*<0.001 in comparison to CDM group). **g** Relative mRNA expression of clock genes in mPFC at ZT06 of naive (n=7), CDM mice (n=5), naïve 6hSD treated (n=6), 6hSD followed by 24hRS (n=6), ketamine injected at ZT00 and tissue harvested 6h (6hKET; n=5) and 30h (30hKET; n=5) later naïve mice, (one-way ANOVA with Bonferroni post-hoc test: \**P*<0.05, \*\**P*<0.01, \*\*\**P*<0.001 in comparison to the naïve group). **h** Schema summarizing the effects of CDM, 6hSD and ketamine on clock gene expression in mPFC – CDM and SD upregulates *Per* and *Cry* clock suppressors, while ketamine downregulates them. Data are presented as mean ±SEM and the individual data points are depicted. See also Supplementary Fig. 3 and Supplementary Data Table 1.

We have recently shown that the mPFC molecular clock plays an important role in mediating the effects of stress and ketamine on depression-like behavior (20). To further understand the antidepressant mechanism of SD, we examined the SD effects on the canonical clock gene expression in the mPFC and compared it to the effects of ketamine. Mice subjected to the CDM paradigm (with induced depressive-like phenotype) were either injected with ketamine or saline at ZT00 or treated with 6h SD (starting at ZT00) and brain tissues were harvested at 8h intervals from ZT06 (the end of the SD protocol) and analyzed with qPCR (Fig. 3a). Identical to the distinct behavioral effects, SD and ketamine also exerted differential impact on mPFC clock gene expression (Figs. 3e, f, Supplementary Fig. S3d). Ketamine reduced the oscillating expression of negative clock genes *Per* and *Cry* in a ZT-specific manner with concomitant significant decrease of the mesor (Per2, Cry1, and Cry2) and amplitude (Per2 and Cry1). In contrast, ketamine upregulated the levels of expression (increased mesor) of the positive regulator *RORɑ*. Meanwhile, SD resulted in ZT-specific elevated oscillating expression of *Per* and *Cry* (increased mesor of *Per2* and *Cry1,* and increased amplitude of *Per2*). Notably, the negative regulator *Rev-erbɑ* expression was suppressed at ZT06 immediately following SD administration, at the point where an antidepressant effect was observed. Thereafter, *Reverbɑ* expression was elevated (showing increased mesor), coinciding with the loss of antidepressant effect following RS 24h after SD (Figs. 3b, c, e, f, Supplementary Fig. S3d).

Moreover, SD and ketamine also had distinct effects on the canonical clock gene expression in the master clock in SCN and the rhythmic behavior of the CDM mice. In contrast to ketamine, 6h of acute SD caused upregulation of *Per1*, *Per2,* and *Bmal1* expression in the SCN at ZT06 and produced a significant 1h phase shift of the locomotor activity onset of the CDM mice at constant darkness conditions (Supplementary Figs. S3a-c).

Next, we tested the effects of 6h SD or ketamine on the mPFC molecular clock and depression-like behavior of naïve mice. Remarkably, SD transiently enhanced *Bmal1* levels and the expression of the negative clock genes *Per1*, *Per2* and *Cry1* in the mPFC of naïve mice. Likewise, repeated swim-stress in the CDM protocol upregulated the expression of several clock suppressors, including *Per1*, *Per2*, *Cry2,* and *Rev-erbɑ,* leading to increased mesor of *Per2* and *Cry2*, and enhanced amplitude of *Rev-erbɑ* (Figs. 3e-g, Supplementary Fig. S3d). Acute ketamine administration on naïve mice at ZT00 resulted in downregulation of *Per1, Cry1,* and *Cry2* expression in mPFC (Fig. 3g). While SD reduced the immobility in both FST and TST immediately after the treatment, the following RS period of 24h led to enhanced depression-like behavior. In contrast, ketamine produced sustained antidepressant effects, also in naïve mice (Fig. 3d).

In general, CDM and SD appear to exert similar potentiating effects on negative circadian genes of the mPFC molecular clockwork, leading to elevated depression-like phenotype in the long term. In contrast, ketamine elicits counter modulatory action on the clock suppressors, resulting in a longer-lasting antidepressant effect (Fig. 3h).

### mPFC *Bmal1* knockout causes day-night specific sleep changes, increases sleep fragmentation, and dysregulates slow wave activity

To examine the importance of the mPFC molecular clock in sleep regulation and in the antidepressant effects of SD, we disrupted *Bmal1* expression in mPFC excitatory glutamatergic neurons, using AAV-mediated expression of Cre-recombinase under the control of the calcium/calmodulin-dependent protein kinase II alpha (CAMK2a) promoter in *Bmal1*flox mice (20). To assess the *Bmal1*KO-induced changes in sleep, following the mPFC injection with Cre or control EGFP AAV, mice underwent ECoG and EMG electrode implantation (Fig. 4a). Our analyses showed that *Bmal1* mRNA levels were successfully downregulated by around 50% in the mPFC of CaMK2a-Cre injected mice, aligning with the relative proportion of CaMK2a-expressing cells in this area (Supplementary Figs. S4a, b), indicating the efficiency of our viral KO strategy. CaMK2a-*Bmal1*KO mice exhibited ZT-specific changes to sleep architecture, spending less time awake in the active dark period at the expense of more time in SWS and REM sleep (Fig. 4b and Supplementary Fig. S4c). In all ZT periods, *Bmal1*KO mice exhibited significant sleep fragmentation, frequently switching between wake and SWS, compared to controls (Fig. 4c and Supplementary Fig. S4d). This fragmentation was also reflected in significantly shorter bouts of wake during the dark period and shorter bouts of SWS across the light period (Figs 4d, e and Supplementary Fig. S4e), as well as in the increased number of REM episodes in the active period (Supplementary Fig. S4f). SWA activity in *Bmal1*KO mice was significantly elevated at several ZT points compared to controls (Fig. 4f) and exhibited no rhythmic changes between the light and dark period (Fig. 4g, Supplementary Fig. S4g). After 6h SD, mPFC *Bmal1*KO mice showed no significant rebound in either SWA or SWS time (Fig. 4h and Supplementary Figs. S4h, i).

**Fig. 4:**
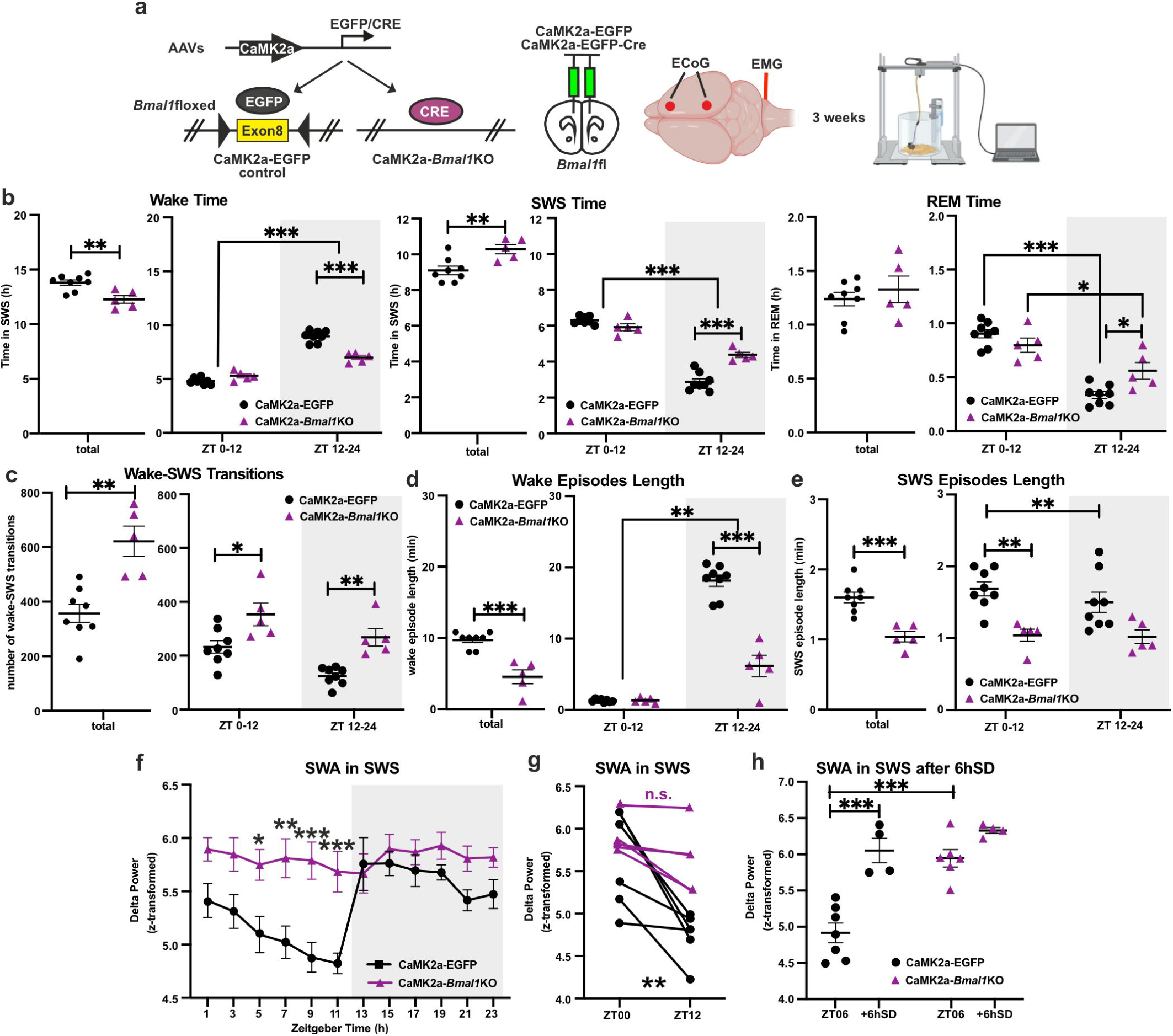
mPFC *Bmal1* knockout causes day-night specific sleep changes, increases sleep fragmentation and dysregulates slow wave activity. **a** Experimental strategy used for region- and cell type-specific virally induced deletion of *Bmal1,* the time course of the AAV microinjections and ECoG/EMG implantation into mPFC and sleep recordings. **b-e** Sleep architecture measures of control CaMK2a-EGFP (n=8) and mPFC CaMK2a-*Bmal1*KO (n=5) mice (left plots - two-tailed Student’s t-test and right plots – repeated measures two-way ANOVA with Bonferroni post-hoc test; \**P*<0.05, \*\**P*<0.01, \*\*\**P*<0.001): **b** Time spent in wake, SWS and REM sleep over the total 24h (left plots) and across 12h light (ZT00-12) and dark (ZT12-24) periods (right plots). **c** Number of wake-SWS transitions over the total 24h (left) and across 12h light (ZT00-12) and dark (ZT12-24) cycles (right). **d-e** Mean duration of spontaneous wake (**d**) and SWS (**e**) episodes for the whole 24h (left) and across 12h light (ZT00-12) and dark (ZT12-24) phases (right). **f** SWA during SWS per 2h episodes across 12h:12h LD conditions of control CaMK2a-EGFP (n=7) and mPFC CaMK2a-*Bmal1*KO (n=5) mice (repeated measures mixed-effects model two-way ANOVA with Bonferroni post-hoc test: \**P*<0.05, \*\**P*<0.01, \*\*\**P*<0.001). **g** SWA in SWS at the start (ZT00) and at the end (ZT12) of the sleep period (repeated measures two-way ANOVA with Bonferroni post-hoc test: \*\**P*<0.001). **h** SWA during SWS at ZT06 at baseline (CaMK2a-EGFP n=7; CaMK2a-*Bmal1*KO n=6) and after 6hSD (n=4 per group) (repeated measures two-way ANOVA with Bonferroni post-hoc test: \*\*\**P*<0.001). Data are presented as mean ±SEM and the individual data points are depicted. See also Supplementary Fig. S4 and Supplementary Data Table 1. Some of the sketches were made with biorender.com

### Antidepressant effects of SD are dependent on mPFC clock function and can be suppressed by REV-ERB agonism

Previously, we have shown that the antidepressant effects of ketamine are dependent on a functional mPFC molecular clock (20). Utilizing the cell- and region-specific *Bmal1*KO, we then aimed to test whether the effects of SD on behavior are also mediated by clock function. CaMK2a-*Bmal1*KO was induced as described above (Supplementary Fig. S4a, b), and the mice were subjected to the CDM paradigm 1 week before being treated with 6h SD. FST and TST were conducted at ZT06, immediately following SD administration (Fig. 5a). While EGFP-injected controls exhibited increased immobility after CDM stress protocol and an antidepressant-like effect after SD, the CaMK2a-*Bmal1*KO mice showed no change in immobility after CDM and subsequent SD, neither in FST nor in TST (Figs. 5b, c).

**Fig. 5:**
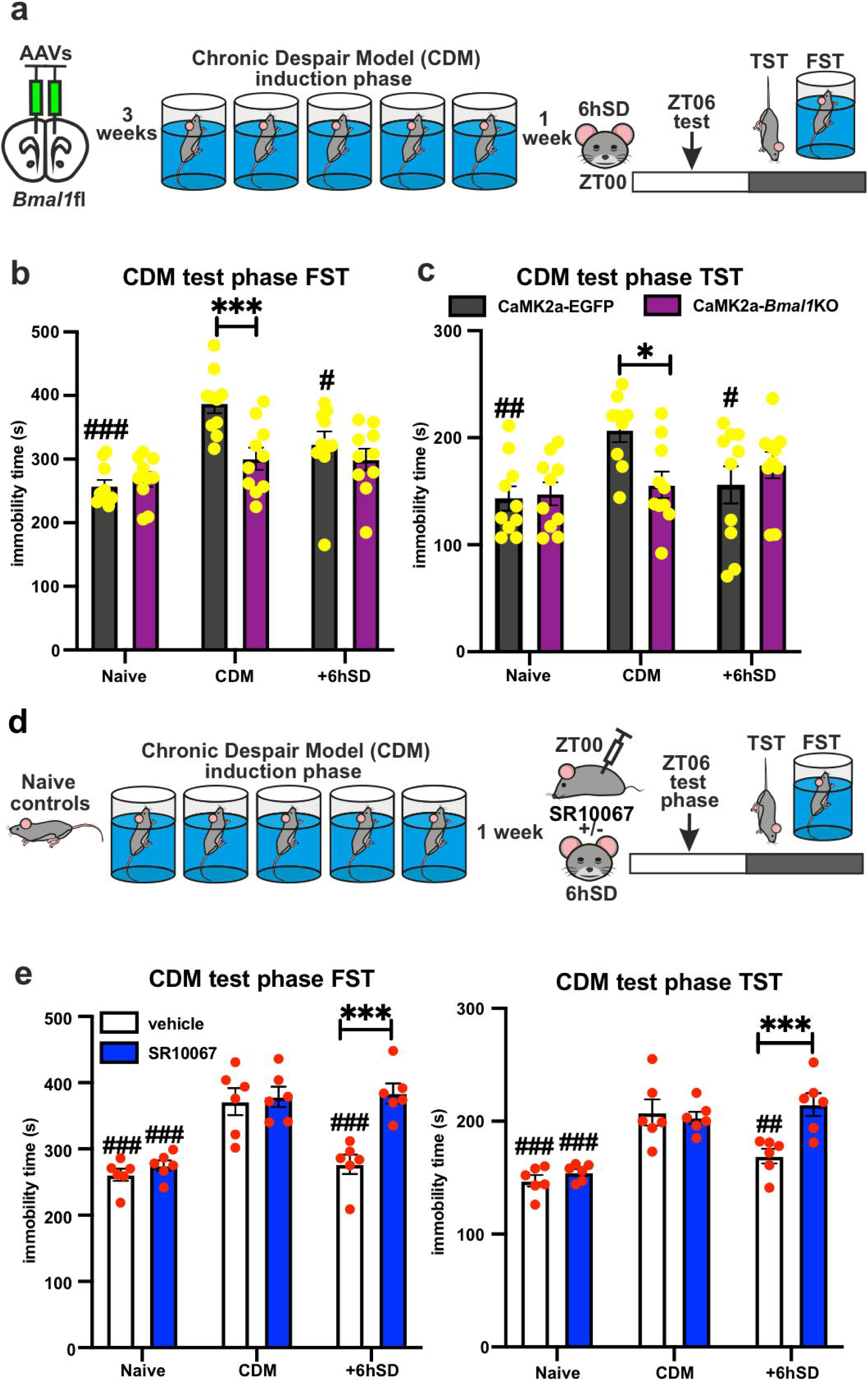
Antidepressant effects of SD are dependent on mPFC clock function and can be suppressed by REV-ERB agonism. **a** Experimental strategy: AAV-induced mPFC CaMK2a-*Bmal1*KO, followed after 3 weeks by CDM paradigm; 1 week later the mice were subjected to 6h SD (ZT00-ZT06) and test phase (TST or FST) at ZT06. **b** Immobility time during FST of CDM day1 (naïve), test phase (CDM) and after 6hSD (n=10 mice per group, repeated measures two-way ANOVA with Bonferroni post-hoc test: \*\*\**P*<0.001 and #*P*<0.05, ##*P*<0.01 vs CDM). **c** Immobility time of TST of naïve mice, CDM mice and CDM after 6hSD (n=10 mice per group, repeated measures two-way ANOVA with Bonferroni post-hoc test: \**P*<0.05 and #*P*<0.05; ##*P*<0.01 vs CDM). **d** Experimental strategy: WT naïve mice injected (at ZT00) with vehicle/SR10067 (30mg/kg) or CDM mice vehicle/SR10067 (30mg/kg) injected (ZT00) and subjected to 6h SD (ZT00-ZT06) were behaviorally tested (at ZT06). **e** Immobility time during test phase FST (left) and TST (right) (n=6 mice per group, two-way ANOVA with Bonferroni post-hoc test: \*\*\**P*<0.001; ##*P*<0.01, ###*P*<0.001 vs CDM). Data are presented as mean ±SEM and the individual data points are depicted. See also Supplementary Data Table 1.

To investigate further the importance of the circadian clock for the antidepressant action of SD, we tested the effect of dual REV-ERBα/β agonist SR10067 (20, 36) on SD administration. WT mice were subjected to the CDM paradigm before being i.p. administered SR10067 (30mg/kg)/vehicle at the beginning of the SD protocol, followed by TST and FST at the end (Fig. 5d). The antidepressant effect observed in vehicle-treated animals was absent in SR10067-injected mice in both the FST and TST. SR10067 alone had no significant effect on the depression-like behavior of naïve and CDM mice (Fig. 5e).

These data demonstrate that the antidepressant effects of SD are dependent on the functional mPFC molecular clock and can be inhibited by pharmacological potentiation of the negative clock regulator REV-ERB.

### mPFC *Bmal1*KO inhibits *Homer1a*-mediated behavioral changes after SD and RS

Next, we investigated the impact of *Bmal1*KO in CaMK2a neurons (Supplementary Fig. S4b) on the mPFC molecular clock functioning and SD effects in naïve nonstressed mice. After the viral induction of the KO, mice were subjected to 6h SD with immediately behavioral testing following SD or after 24h RS at ZT06 (Fig. 6a). In naive EGFP expressing control mice, immobility time was reduced in the FST after SD, and was significantly elevated in comparison to baseline after 24h, suggesting a pro-depressive effect after RS. In contrast, no significant alterations were observed in CaMK2a-*Bmal1*KO mice (Fig. 6b left plot). Likewise, there was no change of TST immobility time after SD nor RS in CaMK2a-*Bmal1*KO mice. Though, SD did not significantly affect the behavior of the EGFP controls in TST, following RS still produced a pro-depressive effect (Fig. 6b right plot). Our qRT-PCR mRNA analyses showed that SD resulted in elevated expression of *Per1, Per2,* and *Cry1* in the mPFC of control mice, while no such effects on the clock suppressors expression were observed in CaMK2a-*Bmal1*KO animals (Fig. 6c). Interestingly, the expression of the positive regulator *RORɑ* was significantly elevated in *Bmal1*KO compared to control mice at baseline, and this expression was reduced after SD to a similar level as seen in controls. Crucially, SD-treated *Bmal1*KO animals exhibited no change in *Homer1a* expression after SD. In contrast, EGFP controls showed a significant Homer1a increase, which has been previously linked to antidepressant effect (19–21, 47).

**Fig. 6:**
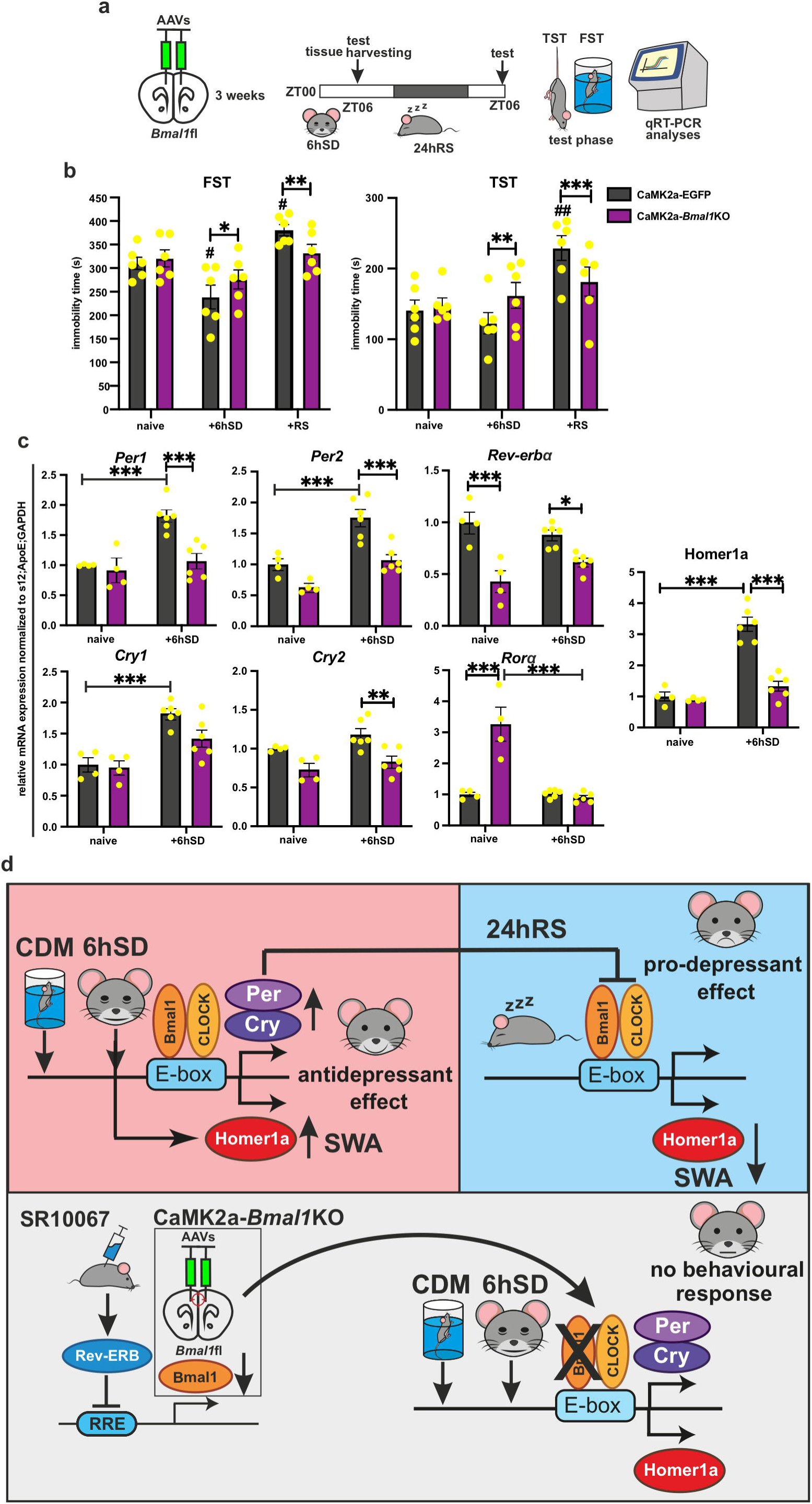
mPFC *Bmal1*KO inhibits *Homer1a*-mediated behavioral changes after SD and RS. **a** Experimental strategy: AAV-induced mPFC CaMK2a-*Bmal1*KO, followed after 3 weeks by 6hSD and 24hRS, TST or FST tested, or tissue harvested at ZT06. **b** Immobility time in FST of naïve mPFC CaMK2a-EGFP/*Bmal1*KO mice at baseline, after 6hSD and following 24hRS (n=6 mice per group, two-way ANOVA with Bonferroni post-hoc test: \**P*<0.05, \*\**P*<0.01, \*\*\**P*<0.001 and #*P*<0.05 vs naive). Immobility time in TST of naïve mPFC CaMK2a-EGFP/*Bmal1*KO mice at baseline, after 6hSD and following 24hRS (n=6 mice per group, two-way ANOVA with Bonferroni post-hoc test: \*\**P*<0.01, \*\*\**P*<0.001 and ##*P*<0.01 vs naive). **c** Relative mRNA expression of clock genes and Homer1a at ZT06 in mPFC of naïve CaMK2a-EGFP/*Bmal1*KO mice at baseline (n=4) and after 6hSD (n=6) (two-way ANOVA with Bonferroni post-hoc test: \**P*<0.05, \*\**P*<0.01, \*\*\**P*<0.001). **d** Model of the circadian clock role in the SD-mediated effects on depression-like behavior: 6h SD potentiates clock suppressor genes (*Per* and *Cry*), temporary increases Homer1a and SWA, leading to rapid antidepressant effect (upper left box). Enhanced negative clock loop inhibits the BMAL1-mediated transcriptional regulation of Homer1a, decreasing SWA and causing pro-depressant effect and/or relapse after 24h RS (upper right box). Pharmacological activation of the clock negative regulator REV-ERB and/or direct *Bmal1*KO in mPFC CaMK2a excitatory neurons leads to decreased/ablated *Bmal1* expression, that blocks the Homer1a and *Per/Cry* modulation by repeated swim-stress (CDM) and SD and inhibits the behavioral response. Data are presented as mean ±SEM and the individual data points are depicted. See also Supplementary Data Table 1.

## DISCUSSION

The rapid antidepressant effect of therapeutic SD has been linked with enhanced glutamatergic signaling and increased SWA during SWS; the positive effect is then lost after RS and associated synaptic downscaling (16, 17, 19, 34). Accumulating evidence indicates potential crosstalk between circadian and homeostatic processes after sleep loss (3, 6, 20, 27, 49–54), underscoring the importance of identifying specific genes and pathways directly involved in the rapid response to SD. Using a stress-based model for depression-like phenotype and region-specific clock modulation, we demonstrate the role of the mPFC molecular clock in regulating mechanisms of sleep architecture and homeostasis (Fig. 4) and mediating the effects of acute SD on behavior (Figs. 5, 6). In addition, our data provide molecular and cellular mechanistic evidence (Fig. 2) to support the integrated synaptic plasticity model of therapeutic SD (7, 31, 33, 34, 49). We propose a potential link between the circadian clockwork and the homeostatic mechanisms associated with the plasticity protein Homer1a and the transient therapeutic action of SD.

Chronic stress is a direct disruptor of sleep and a prevalent risk factor for MDD; in turn, sleep disturbance is considered both a contributing factor to and an important common symptom of MDD (30). To produce a depression-like phenotype in mice, we used the CDM model, in which repetitive swim stress induces a long-lasting increase in immobility time in the classical FST and TST (Fig. 3), alongside reduced reward- and motivation-directed behavior towards sucrose seeking and consumption (as automatically measured in IntelliCage). This paradigm is designed to simulate mild and predictable, but inescapable stress (19, 21, 37–41). We found that CDM alters sleep architecture by fragmenting sleep, indicated by more frequent wake-SWS transitions, shorter bouts of SWS and wake, and an increased number of REM sleep episodes (Fig. 1). Sleep fragmentation is a common phenotype in various animal models of stress-induced depression and MDD patients (30, 55–57). In contrast with several studies in rodents and humans reporting increased REM sleep in depression (30, 55–57), CDM mice exhibited a greater number of REM episodes without an overall change in total REM sleep duration.

Reduction of SWS time has been reported in MDD patients, along with reduced early-night amplitude and altered distribution of associated SWA (30, 57–60). Decreased SWA has been described in preclinical studies after social defeat, chronic swim, and restraint stress (61); however, the time spent in SWS is increased in these models, which may be due to the multi-phasic nature of sleep in mice compared to humans. Consistent with these reports, CDM mice exhibited disruption of the SWA rhythm accompanied by prolonged SWS time, suggesting impaired homeostatic regulation of sleep. In human MDD patients, blunted early-night SWA predicts responsiveness to antidepressant ketamine (62), indicating a treatable link between deficiencies in sleep homeostasis and synaptic plasticity regulation, as predicted by the synaptic homeostasis hypothesis (SHY) (33, 44). Indeed, we have previously shown that ketamine elevates markers of synaptic plasticity, including SWA, and alleviates the depressive-like phenotype in CDM mice (19–21).

According to the SHY (33, 44), SWS and delta activity are bi-directionally associated with both the homeostatic drive for sleep (63) and fundamental mechanisms relating to neuronal homeostatic plasticity (32, 33, 43, 44). SHY proposes SWA as a marker for synaptic strength, with delta power increasing as a function of prior wakefulness and decreasing during sleep, reflecting synaptic downscaling. Impaired plasticity processes may therefore contribute to abnormal SWA (16, 17, 32, 33, 43–45, 64). The expression of the scaffolding protein Homer1a and synaptic trafficking of AMPARs have been identified as strong indicators of homeostatic plasticity and are associated with SWA during SWS (45–47). Homer1a regulates activity-induced bi-directional changes in AMPAR surface expression (65), and plays a key role during late-phase long-term potentiation, where it amplifies AMPAR function (65–67). However, it also drives the homeostatic scaling-down of excitatory synapses during sleep (46). Our results, showing decreased Homer1a levels and AMPAR expression in the mPFC of CDM mice (Fig. 2), add to previous evidence, demonstrating a reduction of synaptic plasticity and glutamatergic signaling (decreased Homer1a levels and AMPAR expression) in both patients and animal models of depression (8, 9, 11, 19, 21, 37, 40, 68–70). These changes are further reflected in the reduction of the day-night amplitude of SWA in CDM mice. Supporting the interconnectedness of these results, we found altered day-night oscillations of Homer1a expression accompanied by lower levels of AMPARs at the end of the active (dark) period, followed by a lack of GluA1 synaptic downscaling during the sleep (light) phase in the mPFC of CDM mice. Indeed, Homer1a has been implicated in both the synaptic downscaling of AMPARs during sleep (46) and the increase of synaptic AMPARs during wakefulness (19).

An integrated synaptic plasticity model of therapeutic SD (34)—combining the synaptic plasticity hypothesis of MDD (7, 10, 12, 13, 26) and the SHY of sleep-wake regulation (32, 33, 44, 45)— proposes that prolonged wakefulness during SD increases overall synaptic strength, which is reduced in MDD, and shifts the synaptic plasticity to the optimal level for network function, leading to therapeutic effect. We have previously demonstrated that the rapid antidepressant effects of SD are mediated by both Homer1a induction and enhanced AMPAR activity in the mPFC (19, 21–23). Consistent with the synaptic plasticity model of therapeutic SD proposed by Wolf and colleagues (34), we show that stressed mice exhibit differential responses to SD in terms of SWA, Homer1a, and synaptic AMPAR induction when compared to their naïve counterparts. Naïve mice have higher baseline levels of Homer1a and AMPAR at the beginning of SD (ZT02) (Fig. 2c, d); this leads to the enhanced induction of Homer1a and SWA elevation (Fig. 2f, g), potentially to increased synaptic downscaling, and ultimately to a lack of AMPAR increase in response to SD (Fig. 2h). In contrast, the lower baseline expression of Homer1a and AMPARs in CDM mice (Fig. 2c, d) results in blunted SD responses of both Homer1a and SWA (Fig. 2f, g), leading to reduced synaptic downscaling and a respective SD-induced increase in AMPAR levels (Fig. 2h). Taken together, these data corroborate the integrated synaptic plasticity model for the antidepressant action of SD (34) and suggest a potential molecular mechanism (23).

While the acute stage of SD produces rapid therapeutic action, its antidepressant effects are only transient. This effect is seen in both human patients and CDM mice, and 24h RS even leads to a pro-depressant-like effect in naïve mice (Fig. 3d). SD and ketamine share similar neurobiological mechanisms of action, involving enhanced glutamatergic signaling (19, 21, 23, 71), although ketamine elicits long-lasting antidepressant effects. The common molecular mechanisms underlying their therapeutic effect may rely on the expression of both clock and synaptic plasticity genes (25). Importantly, the interplay between the circadian clock and the sleep-wake cycle has previously been linked to the regulation of the forebrain’s synaptic transcriptome and proteome (50, 54). Thus, circadian clock modulation may explain the differences in the duration of antidepressant effect (6, 24, 72). Indeed, we found that acute SD induced clock gene expression in the SCN and shifted mouse locomotor activity, while ketamine had no such effect (Supplementary Fig. S3a-c). Dysregulated circadian gene expression has been found in several mood-related brain regions of post-mortem MDD patients (73). Moreover, we have recently shown that several circadian genes of the negative clock regulatory loop are upregulated in the mPFC of various models of stress-induced depression-like phenotype, while ketamine treatment leads to rapid and long-lasting normalization of their expression (20). We show here that, in contrast to ketamine, the effect of SD on clock suppressor genes in the mPFC appears to be analogous to stressors, such as the CDM paradigm (Fig. 3e-h). This may lead to an aggravated depression-like phenotype in the long term, as evidenced by the reversal of the antidepressant effects in CDM and a pro-depressant-like effect in non-stressed mice (Figs. 3b-d, Fig. 6d). Likewise, previous studies have shown similar effects of acute SD on *Per* and *Cry* gene expression in different brain regions (51, 52, 74, 75). While acute SD promotes very rapid antidepressant effects, chronic sleep restriction has detrimental effects on mood regulation and is considered a causative factor for depression.

Several clock gene mouse mutants have been shown to exhibit various alterations in sleep amount, architecture, and response to SD (76). However, a significant limitation of these animal models is their lack of cell-type and brain-region specificity, leading to complex and multifaceted behavioral outcomes. The core circadian clock gene *Bmal1* is regarded as an essential component of the functional molecular clock (77). Our data show that the specific mPFC-specific clock disruption in CaMK2a-Bmal1KO mice leads to an attenuated distribution of sleep-wake cycle across the 24-hour period (with a significantly reduced wake/SWS ratio) and sleep fragmentation (Figs. 4b-e, Supplementary Figs. S4c-f). Notably, mPFC CaMK2a-Bmal1KO animals exhibited dysregulated sleep homeostasis. SWA was consistently elevated throughout the sleep period, showing no LD rhythmicity, and SD failed to induce a significant rebound in either SWA or SWS time. The lack of a functional mPFC clock therefore appears to significantly alter the homeostatic regulation of sleep behavior and physiology. While the elevation of SWA may be explained by excessive sleep fragmentation disrupting synaptic downscaling, such an elevation is not seen in CDM mice, which also exhibited frequent sleep-wake transitions. This suggests a direct influence of the mPFC clock on SWA. Interestingly, whole-body *Bmal1*KO mouse mutants—with *Bmal1* ablated in the SCN, leading to central clock disruption—exhibit a comparable sleep phenotype (78). That such a phenotype emerges in our cell- and region-specific knockout, with an intact central clock, demonstrates a previously unrecognized role of the circadian clockwork of mPFC excitatory neurons in the homeostatic regulation of sleep and related physiological mechanisms.

We have recently shown that mPFC CaMK2a-*Bmal1*KO mice exhibited both stress-resilient and ketamine-resistant behavioral changes, suggesting that *Bmal1* expression and a functional mPFC clock are essential factors for the development of a depression-like phenotype and the opposing therapeutic effect of ketamine (20). Corroborating these data, we demonstrate here that mPFC CaMK2a-*Bmal1*KO also blocks both the antidepressant effects of SD and the pro-depressive-like effects after the following 24h of RS in naïve mice. Accordingly, we show that BMAL1KO inhibits the SD-induced upregulation of the clock suppressor genes in the mPFC. While our molecular analyses were performed on bulk mPFC tissue, our use of a cell-type-specific *Bmal1* knockout model targeting CaMK2a-expressing excitatory neurons demonstrates that the molecular clock in these neurons is critical for SD-induced changes in *Per* and *Cry* expression. Future studies employing single-cell resolution approaches, such as RNAscope, will be essential to further dissect the contributions of distinct neuronal and non-neuronal populations to these circadian and behavioral effects.

Homer1a is an important factor in the mechanism of antidepressant action—its induction in the mPFC is necessary for the effects of several therapeutic treatments, including ketamine and SD, thus representing a common pathway of antidepressant mechanism (19, 21, 22, 40). It has been previously demonstrated that Homer1a undergoes bimodal regulation by the transcriptional factor CREB and the circadian clock. Sato et al. discovered that BMAL1 binds to the *Homer1* promoter in the mouse brain and modulates the Homer1a immediate early response (79). We have recently reported that CaMK2a-*Bmal1*KO disrupts the Homer1a induction required for the antidepressant-like effect of ketamine, and also blocks the downregulation of Homer1a in response to CDM stress (19–23). Likewise, we show here that mPFC CaMK2a-*Bmal1*KO inhibits the SD-mediated Homer1a upregulation necessary for its therapeutic action. We propose that BMAL1 interacts with clock suppressors PERs and CRYs, which are increased by acute SD and post-RS, leading to the downregulation of Homer1a expression and consequently to the reversal of the antidepressant-like effects and/or a pro-depressive-like phenotype (Fig. 6d). Previous studies have shown that suppressing negative clock regulators, such as *Per2* or *Cry1, 2*, in the nucleus accumbens can induce antidepressant-like effects, while their deletion—whether in the whole body, neurons, or glial cells—leads to a resilient-like phenotype (80, 81). Similarly, our recent findings demonstrate that *Per2* knockdown in the mPFC produces antidepressant-like effects, further highlighting the critical role of region-specific negative clock regulators in depression (20).

Another method of influencing the molecular clock, particularly the expression of *Bmal1*, is through the indirect modulation of its repressor REV-ERB (82). Our results demonstrate that pharmacological potentiation of the negative clock element REV-ERB by the synthetic agonist SR10067 (36) inhibits the fast antidepressant effects of acute SD. This is consistent with our previous work showing that SR10067 blocks the rapid therapeutic action of ketamine and causes pro-depressant-like effects, each dependent on the downregulation of both BMAL1 and Homer1a (20).

Taken together, our results identify the circadian clockwork in the mPFC CaMK2a excitatory glutamatergic neurons as a key modulator of the homeostatic regulation and consolidation of sleep. Moreover, we demonstrate the role of mPFC molecular clock modulation in mediating the transient effects of SD on depression-like behavior, specifically the enhanced expression of negative clock regulators and a critical role for BMAL1 and its control of Homer1a oscillation. This work increases our understanding of the mechanism of action of SD and highlights the importance of the circadian clock in both the pathophysiology and rapid treatment of depression. Importantly, the therapeutic modulation of the molecular clock represents a novel strategy for the development of more effective treatments and preventive interventions for depression.

## ACKNOWLEDGMENTS

We thank Dr. James M. Wilson and Dr. Bryan Roth for distributing the pENN.AAV.CamKII.HI.GFP-Cre.WPRE.SV40 and pAAV-CaMKIIa-EGFP viruses, Aurelia Ces and the CompOpt core facility of the UPR3212 for the technical support, Dr. Dominique Ciocca and Dr. Sophie Reibel as well as the whole core facility Chronobiotron UMS3415 (Strasbourg, France) for animal care and animal ethical experimentation support. The study was funded by grants from the German Research Council (SE 2666/2-1 and SE 2666/2-3), Medical Research Foundation (FRM) (AJE201912009450), University of Strasbourg Institute of Advance Studies (USIAS) (2020–035), University of Strasbourg ITI NeuroStra and Centre National de la Recherche Scientifique (CNRS UPR3212) to Tsvetan Serchov; and the Région Grand-Est (Fonds Régional de Coopération pour la Recherche, CLueDol project) for CompOpt equipment.

## AUTHOR CONTRIBUTIONS

Conceptualization, T.S., P.B.; Methodology, W.G., D.S., M.B., C.M., M.V., T.S.; Investigation, W.G., D.S., T.S., A.R., C.R., M.V., S.V.; Resources, P.B., C.N.; Writing - Original Draft, W.G., D.S., T.S.; Writing - Review and Editing, W.G., T.S., D.S., C.N, S.V., M.V.; Funding Acquisition, T.S.; Supervision, T.S., P.B., C.N.

## ADDITIONAL INFORMATION

Supplementary information contains: Supplementary Figures S1-S4 and Supplementary Table S1: Excel file reporting statistical results.

