## Supplementary Information for "The mPFC molecular clock mediates the effects of sleep deprivation on depression-like behavior and regulates sleep consolidation and homeostasis"

**SUPPLEMENTARY INFORMATION**
Supplementary information contains:
Supplementary Figures S1-S4
Supplementary Table S1: Excel file reporting statistical results.

**The mPFC molecular clock mediates the effects of sleep deprivation on depression-like behavior and regulates sleep consolidation and homeostasis**

Wilf Gardner[1,2,7], David H. Sarrazin[1,7], Martin Balzinger[1,2], Carole Marchese[1,2], Axelle Ragno[1], Chockalingam Ramanathan[3], Maxime Veleanu[4], Stefan Vestring[4], Claus Normann[4,5], Patrice Bourgin[1,6], Tsvetan Serchov[1,2,4]*

#### Figure S1

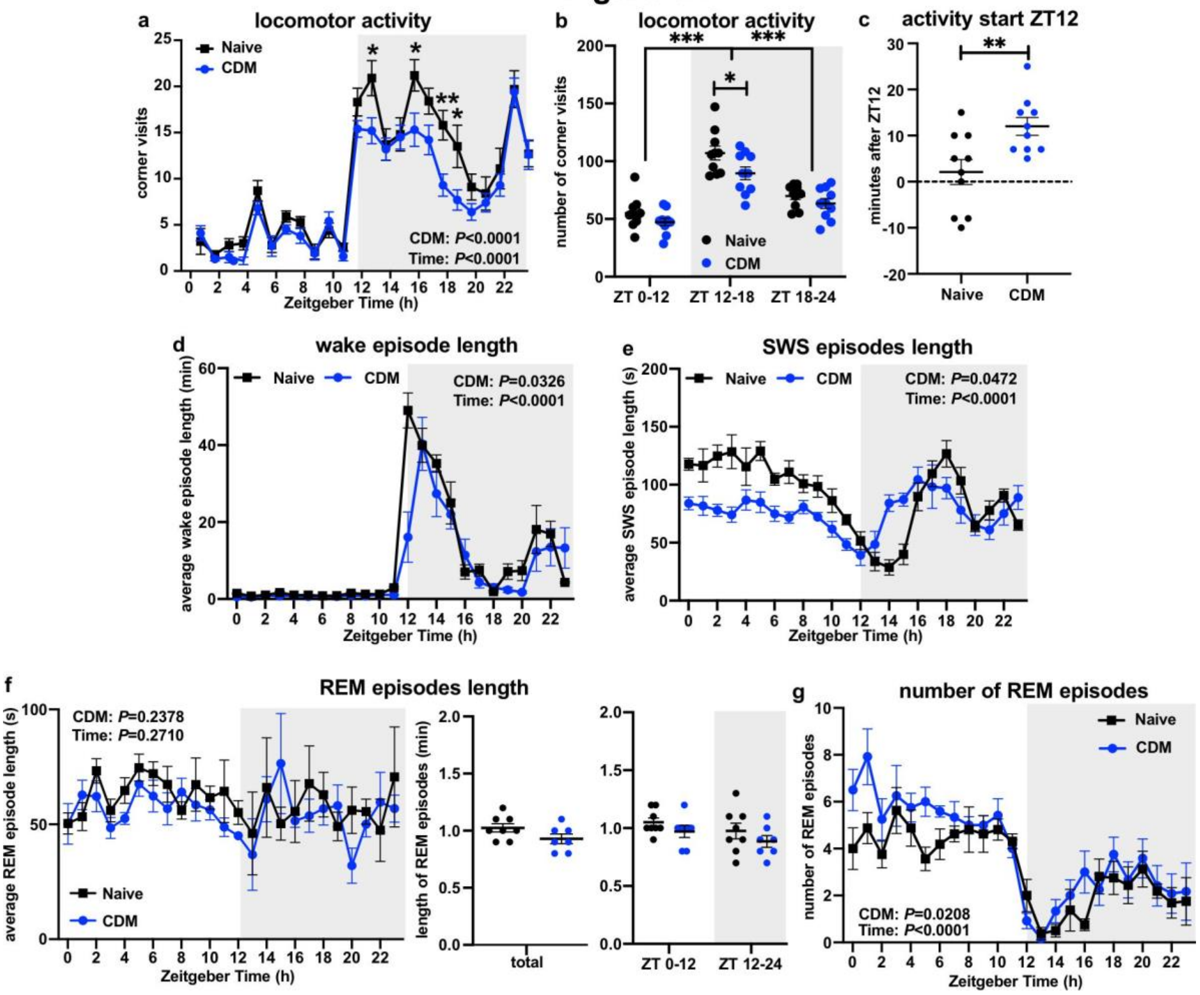


**Figure S1**

**a-b** Locomotor and exploratory activity of naïve (control) and CDM mice (n=10 per group) at 12:12h LD presented as number of corner visits in IntelliCage per 1h periods over 24h (**a**), and for the whole 12h light (ZT00-12), first half (ZT12-18) and second half (ZT18-24) of the dark phase (**b**); repeated measures two-way ANOVA (**a:** *P*-values of the CDM and time effects are displayed inside the graph) with Bonferroni post-hoc test: **P*<0.05, ***P*<0.01, ****P*<0.001;

**c** Mean time onset of the first corner visit (activity) after lights off at ZT12 (n=10 per group, two-tailed Student's t-test: ***P*<0.01).

**d-f** Mean duration of spontaneous wake (**d**), SWS (**e**) and REM (**f**) episodes per 1h periods over 24h and for the whole 24h and across 12h light (ZT00-12) and dark (ZT12-24) cycles (**f** right) of naïve (n=8) and CDM (n=7) mice (repeated measures two-way ANOVA, *P*-values of the CDM and time effects are displayed inside the graphs; two-tailed Student's t-test in **f** middle).

**g** Number of REM episodes per 1h periods over 24h (repeated measures two-way ANOVA: *P*-value of the CDM and time effects are displayed inside the graph).

Data are presented as mean ±SEM and the individual data points are depicted. See also Supplementary Data Table 1.

### Figure S2

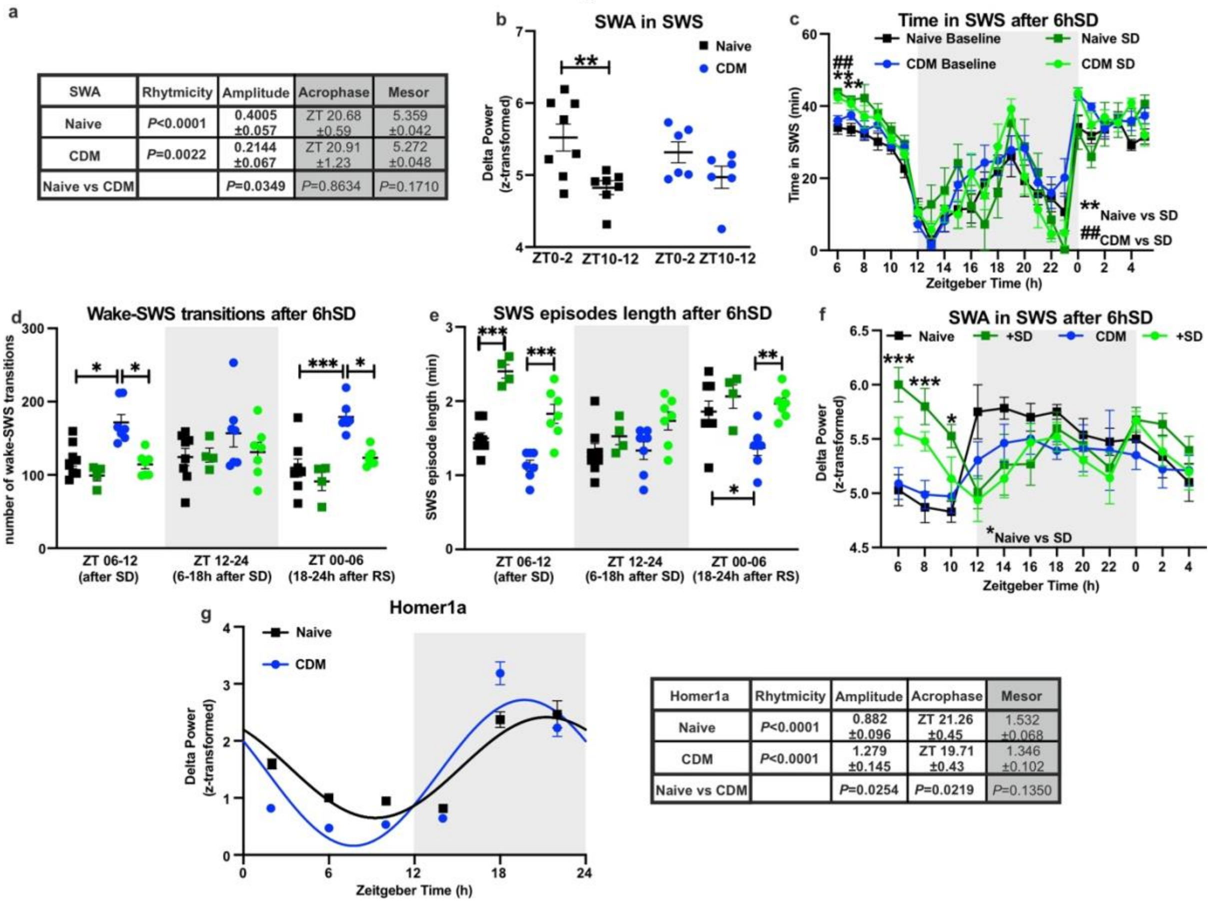


a

| SWA | Rhytmicity | Amplitude | Acrophase | Mesor |
|---|---|---|---|---|
| Naive | *P*<0.0001 | 0.4005 ±0.057 | ZT 20.68 ±0.59 | 5.359 ±0.042 |
| CDM | *P*=0.0022 | 0.2144 ±0.067 | ZT 20.91 ±1.23 | 5.272 ±0.048 |
| Naive vs CDM | | *P*=0.0349 | *P*=0.8634 | *P*=0.1710 |

g

| Homer1a | Rhytmicity | Amplitude | Acrophase | Mesor |
|---|---|---|---|---|
| Naive | *P*<0.0001 | 0.882 ±0.096 | ZT 21.26 ±0.45 | 1.532 ±0.068 |
| CDM | *P*<0.0001 | 1.279 ±0.145 | ZT 19.71 ±0.43 | 1.346 ±0.102 |
| Naive vs CDM | | *P*=0.0254 | *P*=0.0219 | *P*=0.1350 |

**Figure S2**

**a** Comparison of data and statistical significance: rhythmicity, amplitude, acrophase and mesor (naive vs CDM, determined via nonlinear regression sine wave fit and cosinor analyses; extra sum of squares F test) for slow wave activity (SWA) (see also Fig. 1a)

**b** SWA presented as delta power (0.5-4Hz) of ECoG signal recorded during SWS at start (average ZT00-02) and end (average ZT10-12) of the sleep period of naïve (n=8) and CDM (n=6) mice (repeated measures two-way ANOVA with Bonferroni post-hoc test: ***P*<0.01).

**c** Time spent in SWS per 1h periods over 24h at baseline and after 6h of acute sleep deprivation (SD) of naïve (n=8) and CDM (n=7) mice (repeated measures two-way ANOVA with Bonferroni post-hoc test: ***P*<0.01 naïve vs SD, $^{\#\#}$*P*<0.01 CDM vs SD).

**d-e** Number of wake-SWS transitions (**d**) and mean duration of spontaneous SWS episodes (**e**) per 1h periods over 24h at baseline and after 6h of acute sleep deprivation (SD) during the first 6h after SD (ZT06-ZT12), the dark/active period 18h after SD (ZT12-ZT24) and following light/sleep period 24h after SD (ZT00-ZT06) of naïve (n=8) and CDM (n=7) mice (repeated measures two-way ANOVA with Bonferroni post-hoc test: **P*<0.05, ***P*<0.01, ****P*<0.001).

**f** Time course of SWA presented as delta power (0.5-4Hz) during SWS of naïve (n=8) and CDM (n=6) mice at baseline and after 6hSD (repeated measures mixed-effects model two-way ANOVA with Bonferroni post-hoc test: **P*<0.05, ****P*<0.001).

**g** Relative mRNA expression of Homer1a in mPFC of naïve (control) and CDM mice every 4h at 12:12h LD condition (n=5) including fitted sine wave (left) and the calculated amplitude, acrophase and mesor of rhythmic oscillation (right) (naïve vs CDM; nonlinear regression sine wave fit and cosinor analyses; extra sum of squares F test) (see also Fig. 2d).

Data are presented as mean ±SEM and the individual data points are depicted. See also Supplementary Data Table 1.

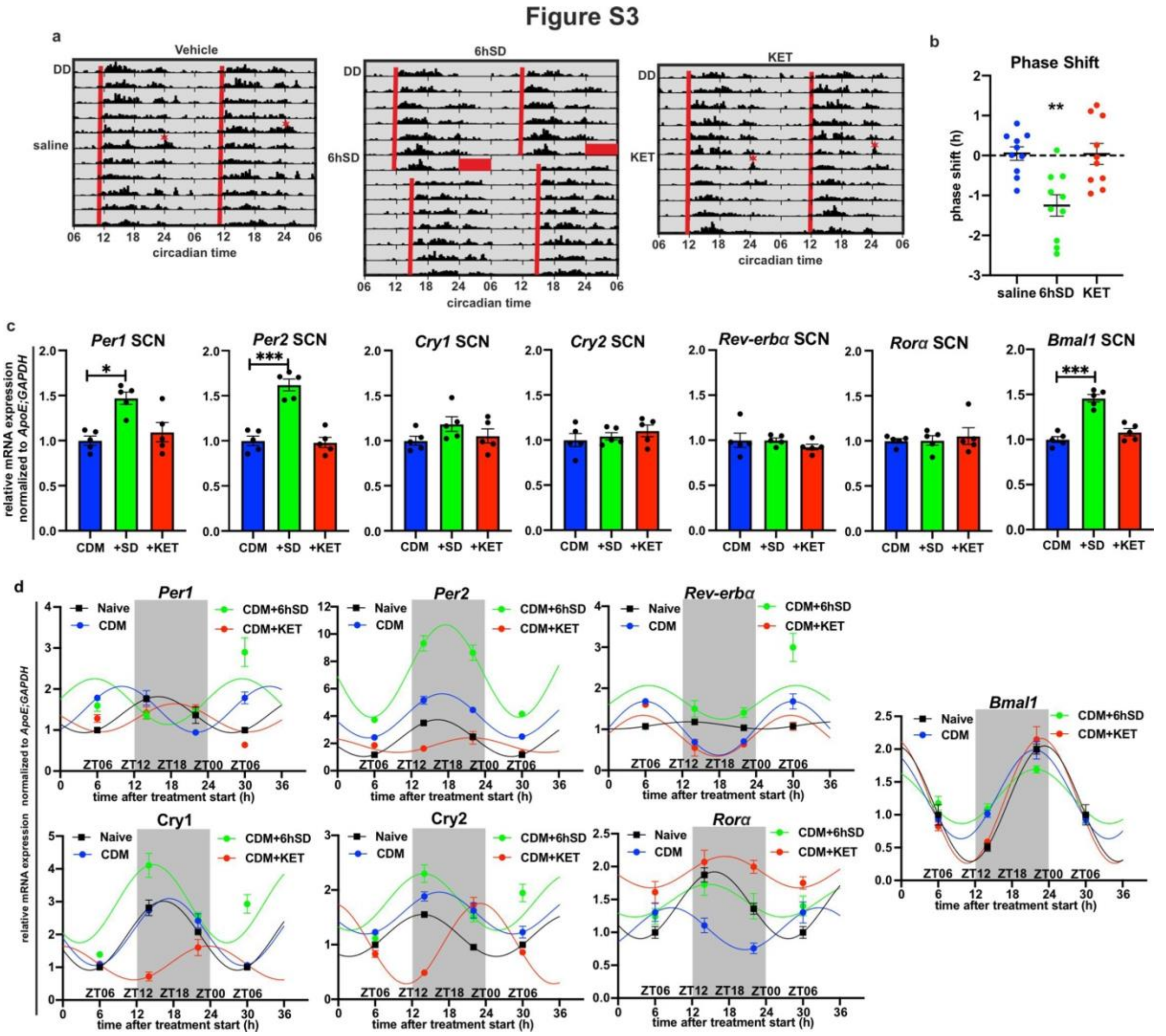


**Figure S3**

**a-b** Representative double plotted actogram (**a**) showing the locomotor activity following saline (vehicle) or 3 mg/kg ketamine (KET) injection at CT00 (red asterisk) or 6h sleep deprivation (6hSD) starting at ZT00 (red box) scored in IntelliCage in freerunning condition (DD), and phase shift quantification (**b**) (n=10 per group; one-way ANOVA with Bonferroni post-hoc test: ***P*<0.01).

**c** Relative mRNA expression of clock genes in SCN at ZT06 of CDM mice, 6h SD treated CDM mice (n=6) and ketamine injected (at ZT00) CDM mice (one-way ANOVA with: **P*<0.05, ****P*<0.001).

**d** Relative mRNA expression of clock genes *Per1, Per2, Cry1, Cry2, Bmal1, Ror*$\alpha$ and *Rev-erb*$\alpha$ normalized to *ApoE* and *GAPDH* in mPFC samples from naïve mice, CDM mice injected at ZT00 with saline (CDM) or ketamine (KET) or 6h SD treated and harvested every 8h from ZT06 (end of SD or 6h after injection) till ZT06 (24h post SD or 30h post injection) including fitted sine waves (n=5 mice per group; nonlinear regression sine wave fit and cosinor analyses) (see also Figure 3e, f).

Data are presented as mean ±SEM and the individual data points are depicted. See also Supplementary Data Table 1.

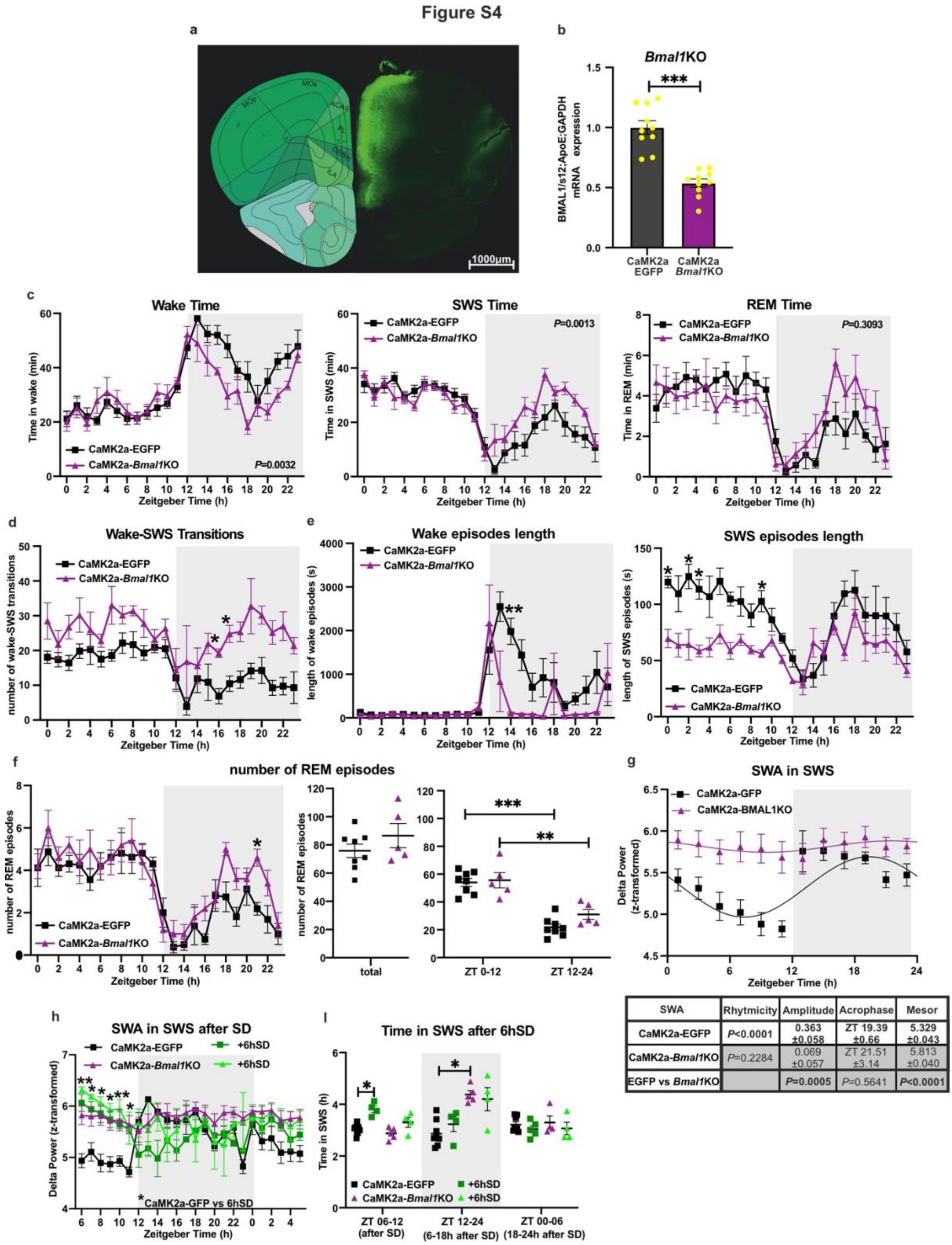


| SWA | Rhytmicity | Amplitude | Acrophase | Mesor |
|---|---|---|---|---|
| CaMK2a-EGFP | *P*<0.0001 | 0.363 ±0.058 | ZT 19.39 ±0.66 | 5.329 ±0.043 |
| CaMK2a-*Bmal1*KO | *P*=0.2284 | 0.069 ±0.057 | ZT 21.51 ±3.14 | 5.813 ±0.040 |
| EGFP vs *Bmal1*KO | | *P*=0.0005 | *P*=0.5641 | *P*<0.0001 |

**Figure S4**

**a** Representative image of viral EGFP/Cre expression in mPFC. Scale bar, 1000µm.

**b** Relative mRNA expression of *Bmal1* in mPFC samples at ZT06 (n=10 per group, two-tailed Student's t-test: ****P*<0.0001).

**c-f** Sleep architecture measures of control CaMK2a-EGFP (n=8) and mPFC CaMK2a-*Bmal1*KO (n=5) mice (repeated measures two-way ANOVA with Bonferroni post-hoc test, *P*-

values of the CDM effect are displayed inside the graph; **f** middle plot: two-tailed Student's t-test; **f** right plot: repeated measures two-way ANOVA with Bonferroni post-hoc test, **$P<0.01$, ***$P<0.001$): **c** Time spent in wake (left), SWS (center) and REM sleep (right) per 1h periods over 24h; **d** Number of wake-SWS transitions per 1h periods over 24h; **e** Mean duration of spontaneous wake (left) and SWS (right) episodes for the whole 24h; **f** Number of REM episodes for the whole 24h and across 12h light (ZT00-12) and dark (ZT12-24) cycles.
**g** SWA during SWS across 12h:12h LD conditions of control CaMK2a-EGFP (n=7) and mPFC CaMK2a-*Bmal1*KO (n=5) mice including fitted sine waves and the comparison of calculated statistical significance for rhythmicity, amplitude, acrophase and mesor of rhythmic oscillation (bellow) (nonlinear regression sine wave fit and cosinor analyses; extra sum of squares F test) (see also Figure 4f).
**h-i** Time spent in SWS per 1h periods over 24h (**g**) and (**h**) during the first 6h after SD (ZT06-ZT12), the dark/active period (ZT12-ZT24) and following 6h of light/sleep period (ZT00-ZT06) of mPFC CaMK2a-EGFP (baseline n=8, SD n=4) and CaMK2a-*Bmal1*KO (baseline n=5, SD n=4) mice (repeated measures two-way ANOVA with Bonferroni post-hoc test: *$P<0.05$, **$P<0.01$, ***$P<0.001$).
Data are presented as mean ±SEM and the individual data points are depicted. See also Supplementary Data Table 1.
